# Creb5 establishes the competence for *Prg4* expression in articular cartilage

**DOI:** 10.1101/2020.08.12.248609

**Authors:** Cheng-Hai Zhang, Yao Gao, Unmesh Jadhav, Han-Hwa Hung, Kristina M Holton, Alan J. Grodzinsky, Ramesh A. Shivdasani, Andrew B. Lassar

**Affiliations:** Department of Biological Chemistry and Molecular Pharmacology, Blavatnik Institute at Harvard Medical School, 240 Longwood Ave., Boston, MA. 02115; Department of Medical Oncology and Center for Functional Cancer Epigenetics, Dana-Farber Cancer Institute, Boston, MA 02215, USA; Departments of Medicine, Brigham & Women’s Hospital and Harvard Medical School, Boston, MA 02215, USA; Department of Biological Engineering, MIT Room NE47-377, 77 Massachusetts Ave, Cambridge, MA 0213; Research Computing, Harvard Medical School, 25 Shattuck Street, Boston, MA 02115; Harvard Stem Cell Institute, Cambridge, MA 02138, USA

**Keywords:** Articular cartilage, chondrocytes, Lubricin, *Prg4*, *Creb5*, Synovial joints

## Abstract

A hallmark of cells comprising the superficial zone of articular cartilage is their expression of lubricin, encoded by the *Prg4* gene, that lubricates the joint and protects against the development of arthritis. Here, we identify Creb5 as a transcription factor that is specifically expressed in superficial zone articular chondrocytes and is required for TGF-β and EGFR signaling to induce *Prg4* expression. Notably, forced expression of Creb5 in chondrocytes derived from the deep zone of the articular cartilage confers the competence for TGF-β and EGFR signals to induce *Prg4* expression. Chromatin-IP and ATAC-Seq analyses have revealed that Creb5 directly binds to two *Prg4* promoter-proximal regulatory elements, that display an open chromatin conformation specifically in superficial zone articular chondrocytes; and which work in combination with a more distal regulatory element to drive induction of *Prg4* by TGF-β. Our results indicate that Creb5 is a critical regulator of *Prg4*/lubricin expression in the articular cartilage.

## Introduction

Osteoarthritis (OA) affects over 30 million US adults (CDC statistics). Therapy for OA is limited to symptom relief and, in severe cases, joint replacement surgery. Interventions that either arrest or reverse the progression of OA are currently unknown. With an eye towards this goal, we have sought to develop a comprehensive understanding of the regulatory network that regulates the differentiation and maintenance of articular cartilage, which plays a central role in maintaining the low-friction environment of the joint space. A hallmark of cells comprising the articular cartilage is their expression of proteoglycans, such as the protein lubricin, encoded by the *Prg4* gene, that lubricates the joint and protects against the development of OA ^1, 2^. *Prg4* is specifically expressed in the superficial-most layer of the articular cartilage, but not by deeper layers of this tissue ^2–7^. Fate mapping studies have established that *Prg4*-expressing cells in embryonic and early post-natal joints constitute a progenitor pool for all regions of the articular cartilage in the adult ^8–10^. These findings are consistent with prior studies indicating that both superficial and deep zones of the articular cartilage (plus other synovial joint tissues) specifically arise from *Gdf5*-expressing cells in the embryo ^11–13^. In both humans and mice lacking *Prg4,* the surface of the articular cartilage becomes damaged and precocious joint failure occurs ^1, 2^. Notably, decreased levels of lubricin have been observed in the synovial fluid following either surgically induced osteoarthritis in sheep ^14^, in human synovial fluid samples from patients with either osteoarthritis (OA) or rheumatoid arthritis (RA) ^15^, and in the menisci from OA patients ^16^. Furthermore, a decrease in *Prg4*/lubricin expression during aging ^17^ correlates with increasing sensitivity of aged knees to cartilage degradation following knee joint destabilization ^18^. Most notably, loss of lubricin (in *Prg4^−/−^* mice) has been noted to result in significantly higher levels of peroxynitrite, superoxide and cleaved Caspase 3 ^19^, which correlates with both increased levels of both whole-joint friction and cellular apoptosis in *Prg4^−/−^* mice compared with either wild-type or *Prg4^+/−^* mice ^20^. Indeed, a hallmark of both aging ^21, 22^ and OA ^23^ is a loss of cells in the superficial zone of the articular cartilage. Conversely, if lubricin protein is either injected directly into the synovial fluid ^24–28^ or over-expressed in the knee joint (via transgene or AAV; ^29, 30^) the articular cartilage tissue is protected from degradation following surgically induced joint-destabilization. Taken together, these findings indicate that *Prg4*/lubricin counters the signaling pathways that lead to cartilage destruction and suggest that identifying a means to induce the sustained expression of *Prg4* in the articular cartilage may attenuate the degradation of articular cartilage observed during either aging or osteoarthritis (OA).

While Foxo ^31^, Nfat ^32^ and Creb1 ^33^ transcription factors (TFs) have been found to modulate *Prg4* expression in articular chondrocytes, the TFs that drive either tissue-specific expression of *Prg4* or region-specific expression of this gene in the superficial zone of articular cartilage have not yet been elucidated. In addition, as *Prg4* is expressed in articular chondrocytes but not in growth plate chondrocytes, we speculated that identification of the TFs that control expression of this gene in the superficial zone of articular cartilage may elucidate how these distinct chondrocyte cell fates are regulated. Several signaling pathways, including Wnt ^11, 34, 35^, TGF-β ^36, 37^ and EGFR ^38^, have all been found necessary to maintain the expression of *Prg4* in the superficial zone of articular cartilage. Interestingly however, injurious mechanical compression ^39^, TGF-β1 exposure ^36^, and shear stress from fluid flow ^33^ can induce *Prg4* expression exclusively in explants taken from the superficial zone of bovine articular cartilage, and not in explants from middle or deep zones. This highly restricted induction implies that superficial zone cells either secrete additional necessary signals or uniquely express a TF(s) that responds to the inductive signals. Here, we identify Creb5 as a TF that is specifically expressed in both bovine and human superficial zone articular cartilage and is critically required to activate *Prg4* expression, in response to TGF-β and EGFR signaling.

## Results

### Creb5, a TF expressed selectively in *Prg4^+^* chondrocytes

In newborn bovine knee joint cartilage, superficial zone chondrocytes (SZCs) expressing *Prg4/*lubricin are readily distinguished from *Prg4/*lubricin-negative deep zone chondrocytes (DZCs, Fig. 1A). As the profound differences between these two tissues likely have a transcriptional basis, we employed RNA-Seq to identify SZC-specific TF genes. Reflecting their common developmental origin ^8–10^, SZCs and DZCs differed by only 320 genes (Supplementary Table 1; false discovery rate, FDR <0.05), including ∼67-fold higher *Prg4* mRNA levels in SZCs, as expected, and >15-fold higher levels of other SZC marker genes such as *Vitrin*, *Epha3*, *Wif1*, and *Thbs4* (Supplementary Table 1; Fig. 1B). Notably, we found that 29 transcriptional regulators were more highly expressed (at least 2-fold) in SZCs than in DCZs; and only one TF (Sall1) was more highly expressed in DZCs (Table 1). The TF gene *Creb5* was the most differentially expressed transcriptional regulator, with ∼25-fold higher levels in SZCs than in DZCs (as determined by RNA-Seq; Fig. 1C), approximately equal to the differential expression of *Prg4* in these tissues. Because TGF-β induces *Prg4* specifically in SZCs *in vitro* ^36^ and maintains *Prg4* expression in articular cartilage *in vivo* ^37^, we assessed mRNA levels in SZCs and DZCs cultured with or without TGF-β2. RT-qPCR revealed ∼10-fold induction of *Prg4* by TGF-β2 only in SZCs, as others have reported ^36^; and a selective 250-fold (*Prg4*) and 40-fold (*Creb5*) greater expression of these transcripts, in response to TGF-β2, in SZCs versus DZCs (Fig. 2A, compare lanes 2 and 4).

**Fig. 1.**
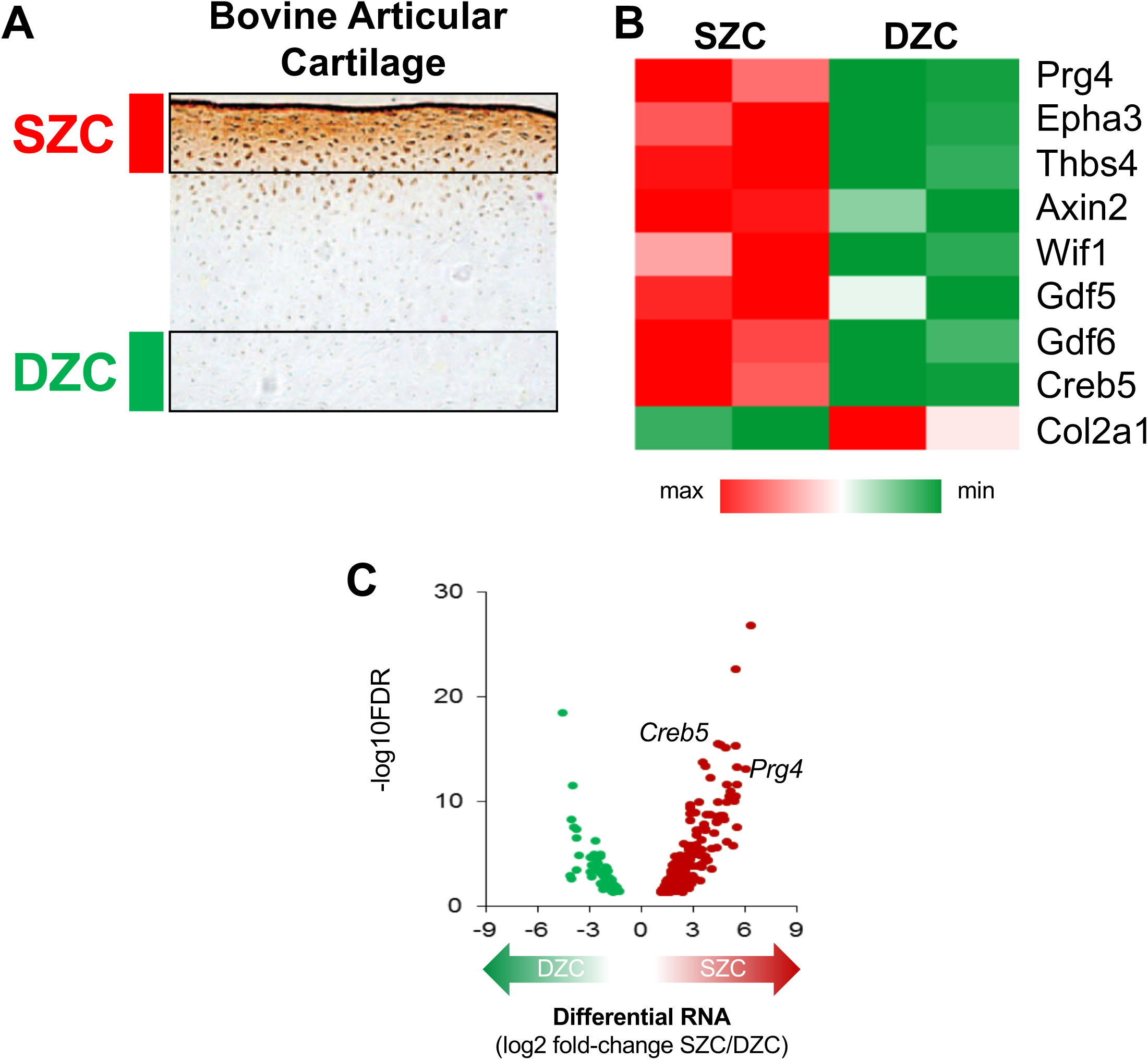
Genes that are preferentially expressed in SZCs versus DZCs. (A) Lubricin (brown) is expressed in superficial zone chondrocytes (SZCs; adjacent to red bar) and not in deep zone chondrocytes (DZCs; adjacent to green bar) in bovine articular cartilage. Image of Lubricin immunostaining taken from ^44^. (B) Relative expression of genes that are differentially expressed in superficial zone bovine articular chondrocytes (SZC) versus deep zone bovine articular chondrocytes (DZC). (C) Volcano plot of differentially expressed genes (DEGs) in SZCs and DZCs. Each dot represents one gene. The red dots represent SZC-specific DEGs, the green dots represent DZC-specific DEGs. Note that the TF gene *Creb5* was as differentially expressed as *Prg4*, with ∼25-fold higher levels in SZCs than in DZCs.

**Fig. 2.**
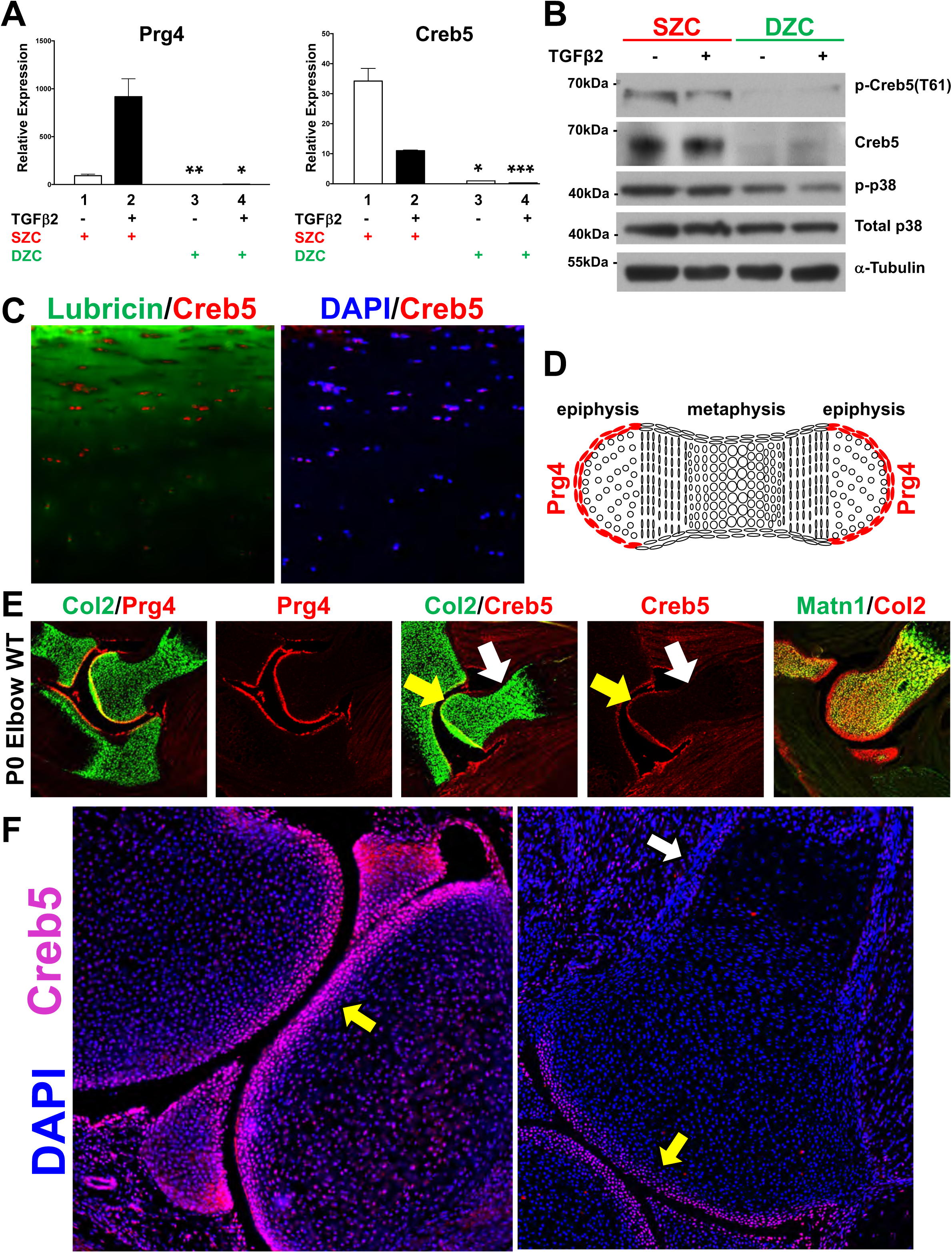
Creb5 is differentially expressed in superficial versus deep zone articular chondrocytes and is specifically expressed in *Prg4^+^* cells in synovial joints. (A) RT-qPCR analysis of *Prg4* and *Creb5* expression in either SZCs or DZCs cultured in either the absence or presence of TGF-β2 (20 ng/ml) for 3 days. Gene expression was assayed by RT-qPCR and normalized to *GAPDH*. Lanes 3 and 4 are compared to lanes 1 and 2, respectively. In both this and subsequent figures: *P<0.05, **P<0.01, ***P<0.001, ND (not detected) and ns (non-significant difference) are indicated and error bar indicates standard error of the mean. Similar results have been obtained in 3 independent biological repeats. (B) Western analysis of proteins in SZCs and DZCs. Similar results have been obtained in 2 independent biological repeats. (C) Immunofluorescent staining for CREB5, Lubricin and DAPI (to visualize nuclei) in adult human femoral head articular cartilage. (D) Schematic of *Prg4* expression in a developing long bone cartilage element. (E) Expression of *Collagen2a1 (Col2), Prg4, Creb5,* and *Matrilin1 (Matn1)* in the elbows of P0 mice, as detected by fluorescent in situ hybridization. (F) Immunofluorescent staining for Creb5 and DAPI (to visualize nuclei) in the knee joint of a P0 mouse. In (E & F), the *Creb5*-expressing articular perichondrium is designated by a yellow arrow; the metaphyseal perichondrium is designated by a white arrow.

**Table 1.**
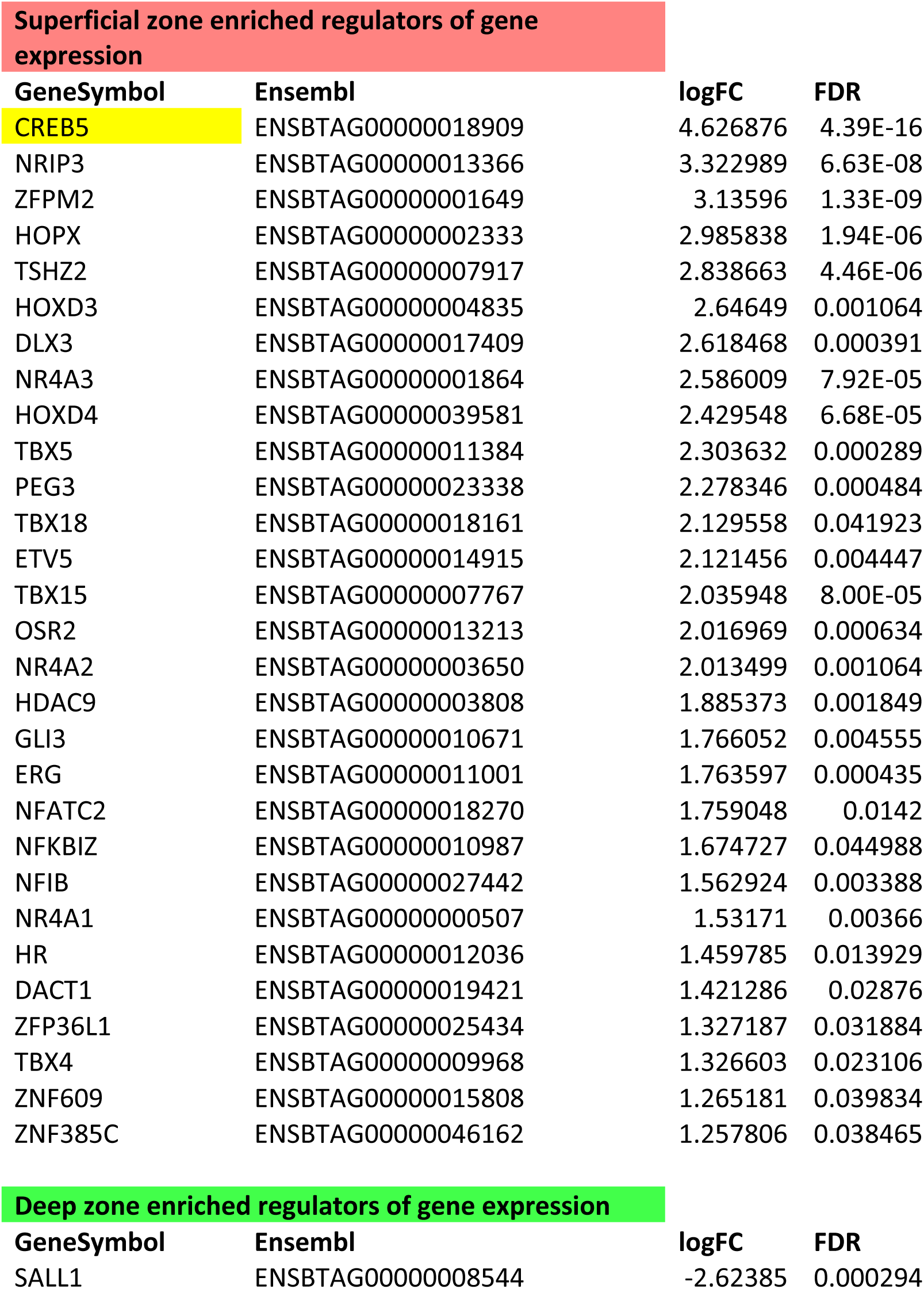
Differentially expressed regulators of gene expression. Transcriptional regulators are listed whose expression differed (by indicated LogFC) between SZC and DZC (false discovery rate, FDR <0.05).

The basic-leucine zipper (bZIP) DNA-binding domain of Creb5 shares high sequence homology with those of Atf2 ^40^ and Atf7 ^41^. In addition, all three TFs carry two conserved N-terminal Threonine/Proline residues (T59 and T61 in Creb5) that are substrates for the P38, Jun N-terminal (JNK), and extracellular signal-regulated (ERK) kinases (reviewed in ^42, 43^). Consistent with selective *Creb5* mRNA expression in SZCs, a ∼65-kDa protein recognized by both Creb5 and phospho-specific Creb5(T61) antibodies is enriched specifically in SZCs (Fig. 2B). Infection of SZCs with lentivirus encoding an shRNA directed against the 3’UTR of *Creb5* substantially diminished these protein levels (Fig. 3C). Of note, phospho-P38 kinase, the active form that can phosphorylate Creb5 on T61, is also more abundant in SZCs than in DZCs (Fig. 2B). Although TGF-β2 diminished the level of *Creb5* mRNA to ∼30% of that in untreated SZCs (Fig. 2A, lanes 1 and 2), this decrease had no appreciable effect on either total or phospho-Creb5 protein levels (Fig. 2B and Fig. 3C). Consistent with the restricted expression of *Creb5* in the superficial zone of bovine articular cartilage, we similarly detected nuclear-localized Creb5 specifically in the lubricin-expressing superficial zone of articular cartilage in adult human femoral head tissue (Fig. 2C)

**Fig. 3.**
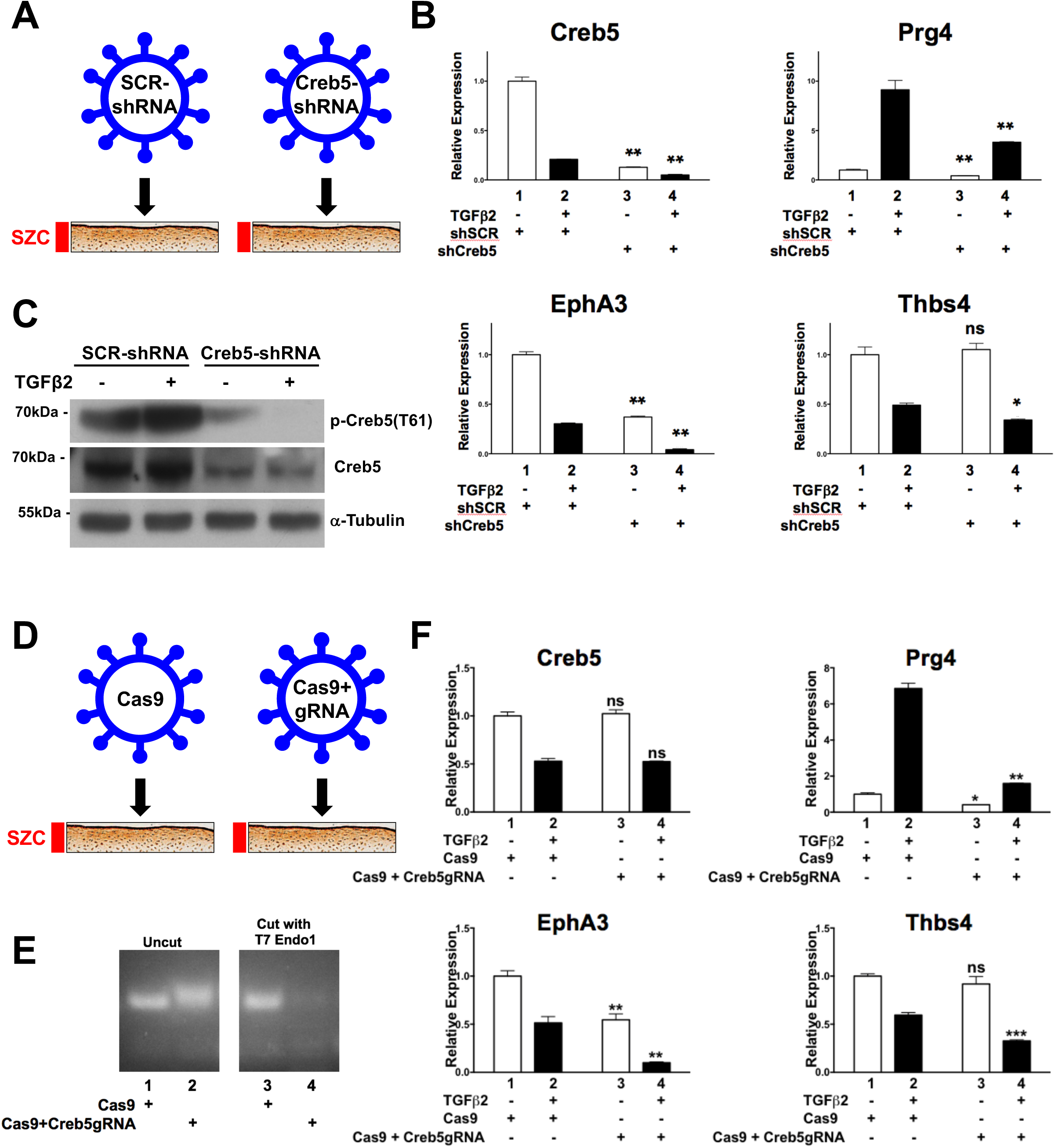
Creb5 function is necessary for TGF-β-dependent induction of *Prg4* in superficial zone articular chondrocytes. (A-C) shRNA-mediated knock-down of Creb5 in superficial zone articular chondrocytes. SZCs were infected with a lentivirus encoding either control scrambled shRNA (shSCR, lanes 1 & 2) or shCreb5 (lanes 3 & 4), after selection in puromycin, the cells were cultured in either the absence or presence of TGF-β2 (20 ng/ml) for 3 days. (B) Gene expression was assayed by RT-qPCR and normalized to *GAPDH*. Lanes 3 and 4 are compared to lanes 1 and 2, respectively. Similar results have been obtained in 3 independent biological repeats. (C) Western analysis of proteins in SZCs recognized by antibodies directed against either phospho-Creb5 (T61), total Creb5, or α-tubulin. Similar results have been obtained in 2 independent biological repeats. (D-F) CRISPR-Cas9 mediated mutation of the DNA binding domain of *Creb5* in SZCs. SZCs were infected with a lentivirus encoding Cas9 alone or Cas9 plus a *Creb5* guide RNA (targeting the DNA binding domain), as indicated. After selection in puromycin, the cells were cultured in either the absence or presence of TGF-β2 (20 ng/ml) for 3 days. (E) T7 Endonuclease 1 assay (which cleaves at mismatches) is displayed for the RT-PCR amplicon encoding the bZIP domain of *Creb5*. (F) Gene expression was assayed by RT-qPCR and normalized to *GAPDH*. Lanes 3 and 4 are compared to lanes 1 and 2, respectively. Similar results have been obtained in 2 independent biological repeats.

*Prg4* is initially expressed in the articular perichondrium, which encases the epiphyses of the developing long bones (depicted schematically in Fig. 2D); and is subsequently expressed in the superficial-most layer of mature articular cartilage, but not by deeper layers of this tissue ^2–7^. We used RNA in situ hybridization to localize *Creb5* transcripts in relation to *Prg4* and *Collagen 2a1 (Col2a1)* in the elbow joints of newborn mice. Consistent with our findings in bovine articular chondrocytes, we detected *Creb5* transcripts in *Prg4*-expressing cells (Fig. 2E). More specifically, *Creb5* expression was restricted to the articular perichondrium (Fig. 2E, yellow arrow), where *Prg4^+^* precursor cells are known to generate articular cartilage ^8–10^, and was absent from perichondrial cells adjacent to the nascent metaphyseal growth plates of developing long bones (Fig. 2E, white arrow). While articular chondrocytes express *Col2a1* but not *Matrilin1 (Matn1)*, epiphyseal chondrocytes (which will later undergo endochondral ossification) express both *Matn1* and *Col2a1* ^10^. Notably, expression of *Creb5* (and *Prg4*) is restricted to the articular (*Col2a1^+^/Matn1^−^*) chondrocytes (Fig. 2E). In developing mouse knees, *Prg4* is expressed in the articular perichondrium, in superficial cells of the prospective meniscus, and in synovial fibroblasts that line the joint cavity ^2, 4, 44^. Creb5 immunocytochemistry on newborn mouse knees indicated that Creb5 protein was specifically expressed and nuclear localized in these very regions (Fig. 2F, yellow arrow designates the articular perichondrium); and was absent from the metaphyseal perichondrium adjacent to the growth plate (Fig. 2F, white arrow). Thus, Creb5 co-localizes precisely with cells that express *Prg4* in newborn bovine and murine joints, and in adult human articular cartilage.

### Creb5 is necessary for induction of *Prg4* by TGF-β

To examine the role of Creb5 in regulating SZC-specific genes in articular chondrocytes, we infected primary bovine SZCs with a control lentivirus (encoding puromycin resistance and carrying a scrambled shRNA) or one engineered to express an shRNA directed against the *Creb5* 3’UTR (Fig. 3A). After selection in puromycin, we cultured the cells for 3 additional days with or without TGF-β2, in ultra-low attachment dishes to induce a round cell shape, which favors chondrogenic differentiation ^45^. The shRNA against the *Creb5* 3’UTR significantly attenuated *Creb5* transcripts (Fig. 3B) and protein (Fig. 3C), and reduced TGF-β2 induction of *Prg4* by 58% (Fig. 3B; compare lanes 2 and 4). Because shRNAs can have off-target effects ^46^, we also infected primary SZCs with lentivirus encoding puromycin-resistance and Cas9 without (control) or with a guide RNA targeting cleavage within the *Creb5* DNA-binding domain (Fig. 3D). After selection in puromycin and culture with or without TGF-β2, T7 Endonuclease 1 assays ^47^ confirmed efficient introduction of indels in the targeted bZIP domain of *Creb5* (Fig. 3E). While 33% of indels (insertion/deletions) induced by CRISPR/Cas9 mutagenesis are predicted to generate mutations within the bZIP domain that still maintain the reading frame for Creb5; the remaining 67% will produce out of frame proteins. However, as the bZIP domain is sufficient for DNA binding by Creb5 ^40^, either mutation within this domain or production of a truncated out of frame fusion protein would be expected to significantly affect the ability of Creb5 to induce target gene expression. Consistent with the notion that Creb5 DNA binding activity is necessary for TGF-β dependent expression of *Prg4*, TGF-β2 induction of *Prg4* was reduced by 77% in SZCs containing indels in the *Creb5* bZIP domain (Fig. 3F, compare lanes 2 and 4). Amongst the handful of other SZC-specific genes whose expression we assayed by RT-qPCR, either shRNA-mediated knock-down of *Creb5* or CRISPR/Cas9-generated indels in the bZIP domain of *Creb5* decreased baseline expression of *Epha3*, but not that of *Thbs4* (Figs. 3B and 3F; compare lanes 1 and 3). Interestingly however, in the presence of TGF-β2, either shRNA-mediated knock-down of *Creb5* or loss of Creb5 DNA interaction decreased the levels of both *Epha3* and *Thbs4* (Figs. 3B and 3F; compare lanes 2 and 4). As TGF-β2 administration reduced *Creb5* transcript levels but not Creb5 protein levels, it is possible that this treatment both destabilizes pre-existing RNAs in SZCs (such as *Creb5, Epha3,* and *Thbs4*) while inducing the expression of *Prg4*, which requires this signal for its expression ^36, 37^. Taken together, these findings suggest that Creb5 is necessary to both maintain expression of some SZC-specific genes (i.e., *Epha3* and *Thb4*) in the presence TGF-β signaling; and is essential for TGF-β signals to induce maximal *Prg4* expression.

### Creb5 confers competence for *Prg4* expression in DZCs

As both TGF-β ^36, 37^ and EGFR ^38^ signaling promote *Prg4* expression in articular cartilage, we asked if these pathways might regulate *Creb5* expression or phosphorylation. Treatment of primary bovine SZCs with TGF-β2, but not the EGFR agonist TGF-α, reduced *Creb5* mRNA (Fig. 4A, lanes 2 and 3). However, in SZCs the combination of TGF-β2 and TGF-α synergistically boosted both *Prg4* expression (Fig. 4A, lane 4) and phospho-Creb5 (T61) levels (Fig. 4B). Because the highly biased expression of Creb5 in SZCs versus DZCs correlates with the restricted ability of TGF-β to induce *Prg4* expression in superficial zone cells (Fig. 2A), we next asked whether Creb5 is the key factor distinguishing the competence for *Prg4* induction in SZCs versus DZCs. To this end, we infected DZCs with a pInducer20 lentivirus ^48^ engineered to express a doxycycline-inducible *Creb5* cDNA appended with 3 carboxy-terminal hemagglutinin epitope tags (iCreb5-HA). Indeed, treatment of iCreb5-HA infected DZCs with doxycycline, TGF-β2 and TGF-α boosted *Prg4* expression ∼44-fold (Fig. 4C, compare lanes 1 and 7), equal to *Prg4* levels in SZCs treated with the same ligands (Fig. 4C, lane 9). In contrast, iCreb5-DZCs treated with TGF-β2 and TGF-α failed to activate *Prg4* in the absence of doxycycline (Fig. 5E, lane 6). Thus, forced expression of Creb5 in DZCs is sufficient to promote the competence for *Prg4* induction by EGFR and TGF-β signals.

**Fig. 4.**
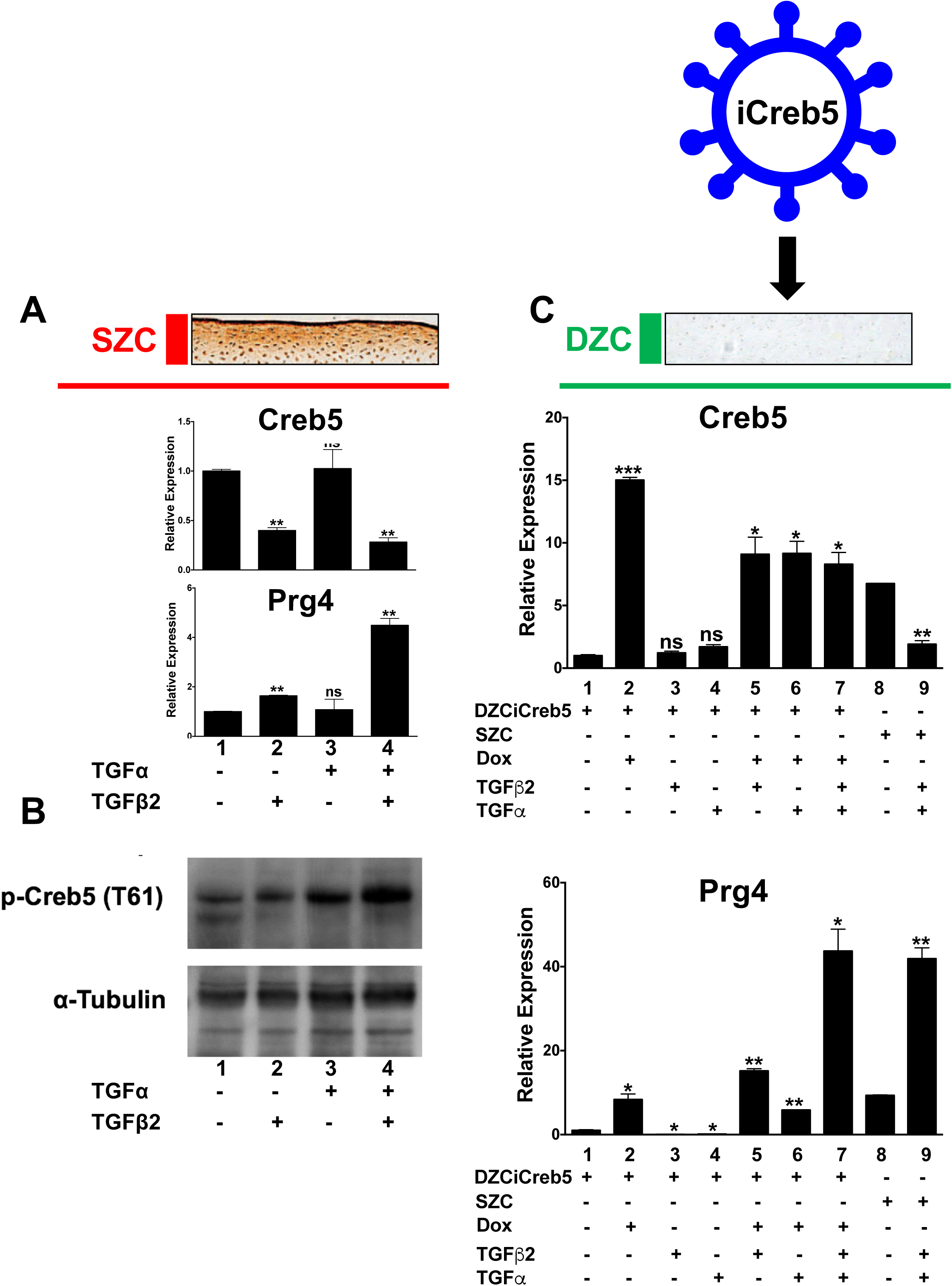
Forced expression of Creb5 in deep zone articular chondrocytes confers competence for EGFR and TGF-β signals to induce expression of *Prg4*. (A & B) EGFR and TGF-β signals synergistically induce expression of *Prg4* in SZCs. SZCs were treated for 2 days with TGF-β2 (20 ng/ml) and TGF-α (100 ng/ml), as indicated. (A) Gene expression was assayed by RT-qPCR and normalized to *GAPDH*. Lanes 2-4 are compared to lane 1. Similar results have been obtained in 3 independent biological repeats. (B) Western analysis of proteins in SZCs. Similar results have been obtained in 2 independent biological repeats. (C) Forced expression of Creb5 in DZCs promotes synergistic induction of *Prg4* by TGF-α and TGF-β2. DZCs were infected with a lentivirus encoding doxycycline-inducible Creb5. After selection in G418, the cells were cultured in either the absence or presence of doxycycline (1μg/ml), TGF-β2 (20 ng/ml) and TGF-α (100 ng/ml), as indicated for 3 days. Gene expression was assayed in both the iCreb5-DZCs (lanes 1-7) and in control SZCs (lanes 8-9) by RT-qPCR and normalized to *GAPDH*. Lanes 2-7 are compared to lanes 1; lane 9 is compared to lane 8. Similar results have been obtained in 3 independent biological repeats.

**Fig. 5.**
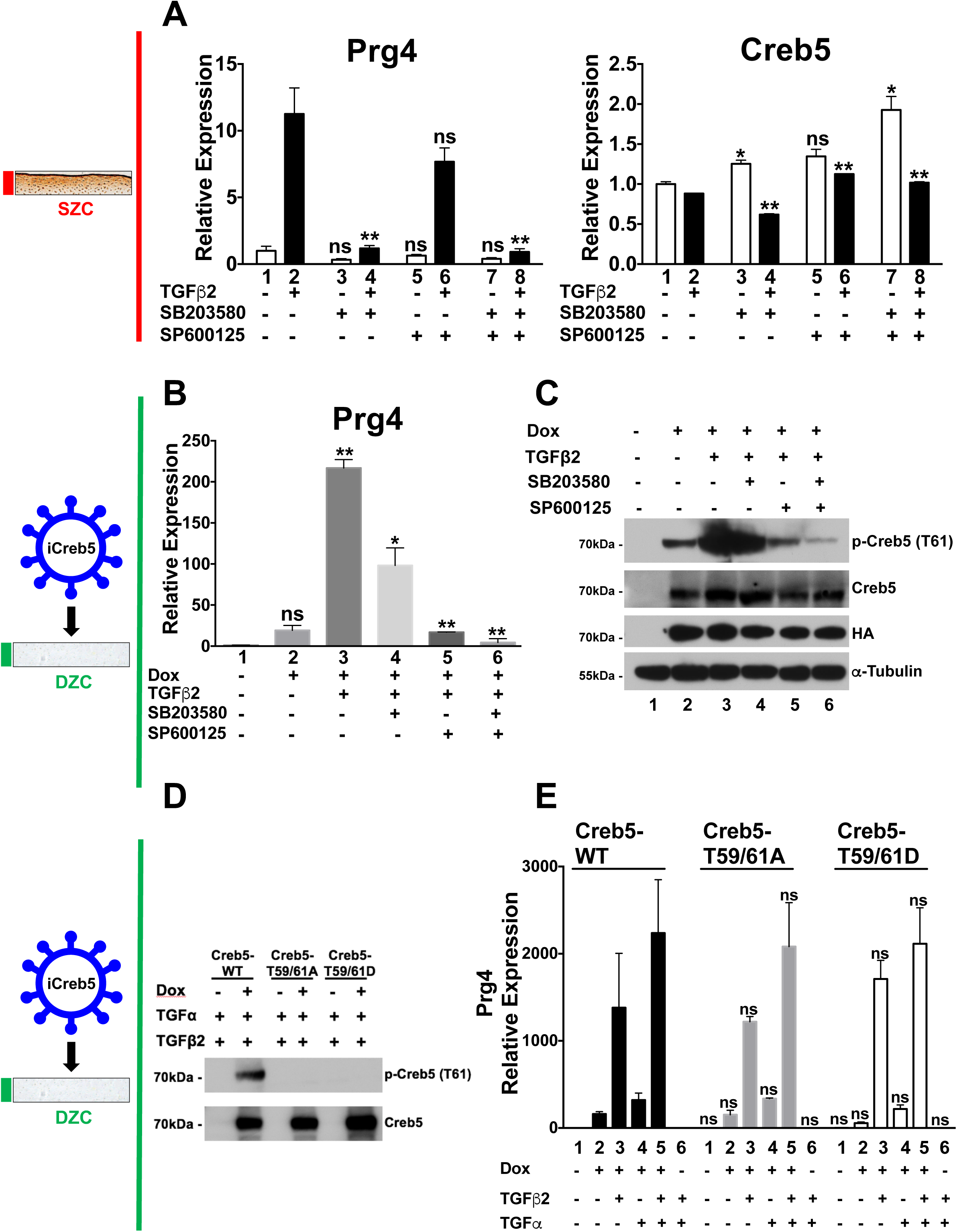
Creb5-dependent induction of Prg4 requires SAPK activity, but neither phosphorylation of T59 nor T61 in Creb5. (A) TGF-β2 induced expression of *Prg4* in superficial zone articular chondrocytes is blocked by inhibition of p38 activity. SZCs were cultured in either the absence or presence of TGF-β2 (20 ng/ml), a p38 inhibitor (SB203580; 10 μM) or a JNK inhibitor (SP600125; 10 μM) as indicated for 2 days. Gene expression was assayed by RT-qPCR and normalized to *GAPDH*. Odd and even lanes are compared to lanes 1 and 2, respectively. Similar results have been obtained in 3 independent biological repeats. (B & C) DZCs were infected with a lentivirus encoding doxycycline-inducible Creb5-HA. After selection in G418, the cells (iCreb5-DZCs) were cultured in either the absence or presence of doxycycline (1μg/ml), TGF-β2 (20 ng/ml), a p38 inhibitor (SB203580; 10 μM) or a JNK inhibitor (SP600125; 10 μM), as indicated for 2 days. (B) Gene expression was assayed by RT-qPCR and normalized to *GAPDH*. Lanes 2-3 are compared to lane 1; lanes 4-6 are compared to lane 3. Similar results have been obtained in 3 independent biological repeats. (C) Western analysis of exogenous Creb5 expression/phosphorylation in iCreb5-DZCs. Similar results have been obtained in 2 independent biological repeats. (D & E) DZCs were infected with a lentivirus encoding either iCreb5-WT-HA; iCreb5-T59,61A-HA; or iCreb5-T59,61D-HA. After selection in G418, the cells were cultured in either the absence or presence of doxycycline (1μg/ml), TGF-β2 (20 ng/ml) and TGF-α (100 ng/ml) as indicated for 3 days. (D) Western analysis of iCreb5-WT/mutant expression in iCreb5-DZCs. Similar results have been obtained in 2 independent biological repeats. (E) Gene expression was assayed by RT-qPCR and normalized to *GAPDH*. Relative level of *Prg4* gene expression driven by the iCreb5-mutant was compared to that induced by iCreb5-WT, under each experimental condition. Similar results have been obtained in 3 independent biological repeats.

In the limbs of newborn mice, expression of both *Creb5* and *Prg4 is* robust in articular chondrocytes and is absent from growth plate chondrocytes (Fig. 2F). To begin to determine whether *Creb5* can confer competence for *Prg4* expression in growth plate-like chondrocytes, we infected either a human chondrosarcoma cell line (SW1353) or an immortalized human costal chondrocyte cell line (C-28/I2;^49^) with lentivirus encoding either EGFP or Creb5. Notably, forced expression of Creb5 conferred competence for TGF-β signaling to induce *PRG4* expression in both cell lines (Supplementary Fig. 1). However, in contrast to deep-zone bovine articular chondrocytes, in which the combination of *Creb5* and TGF-β signaling promotes relatively high-level expression of *Prg4* (approximately equal to 75% of *Gapdh* transcript levels), in the human chondrogenic cell lines, the combination of *Creb5* and TGF-β signaling only induced *PRG4* expression to approximately 0.2% of *GAPDH* transcript levels. Taken together, these findings indicate that *Creb5* can confer competence for *Prg4* induction; and suggest that this TF may work together with other factors in articular chondrocytes to promote high-level expression of *Prg4* in response to TGF-β signals.

### Creb5-dependent induction of *Prg4* requires SAPK activity

Because TGF-β2 and TGF-α synergistically boosted both *Prg4* expression and Creb5 (T61) phosphorylation, we asked whether stress-activated protein kinases (SAPKs) are necessary to induce *Prg4* expression. In SZCs, TGF-β2 induction of *Prg4* was specifically blocked by SB203580, an inhibitor of p38 kinase, but not by SP600125, a JNK antagonist (Fig. 5A); p38 inhibition altered *Creb5* transcript levels only slightly (Fig. 5A). In iCreb5-DZCs, induction of *Prg4* by TGF-β2 was partially blocked by both inhibitors, and completely blocked by the combination (Fig. 5B). Thus, SAPKs are necessary to promote *Prg4* expression by either endogenous or exogenous Creb5 in chondrocytes. In addition, the combination of SAPK inhibitors significantly decreased iCreb5 phosphorylation on T61 (Fig. 5C). To clarify whether SAPK phosphorylation of T59 and T61 on Creb5 is necessary for *Prg4* induction, we mutated both these residues to alanine to block phosphorylation, or to aspartic acid to mimic constitutive phosphorylation. The transcriptional activity of a chimeric protein containing the GAL4 DNA binding domain fused to the N-terminus of Creb5 (GAL4-Creb5-(1-128) was attenuated by simultaneous T59A/T61A mutations, suggesting that phosphorylation of these residues may indeed promote Creb5 transcriptional activity (Supplementary Fig. 2). Interestingly however, a chimeric protein containing full length Creb5 (GAL4-Creb5-(1-508)) drove significantly greater target gene expression than a chimeric protein containing only the N-terminus of Creb5 (GAL4-Creb5-(1-128), suggesting that regions outside the N-terminus of this protein can also drive transcriptional activation (Supplementary Fig. 2). As anticipated, phospho-Creb5 (T61) antibody failed to recognize both mutant forms of Creb5 (Fig. 5D). Surprisingly however, in response to TGF-β2 and/or TGF-α, both mutant iCreb5 forms induced *Prg4* expression to the same level as did wild-type iCreb5 (Fig. 5E). Thus, although SAPKs promote Creb5 T61 phosphorylation, their requirement for *Prg4* induction must depend on phosphorylation either of other proteins, or of Creb5 sites other than T59 and T61.

### Identification of *Prg4* regulatory elements

Because the above findings collectively implicate Creb5 as a key regulator of *Prg4* expression, we sought to understand the basis for its crucial role in driving SZC-specific gene expression. Active enhancers and promoters are both marked by relatively accessible regions of chromatin (reviewed in ^50^). To identify genomic sites with differential chromatin access in SZCs and DZCs, we performed the Assay for Transposase-Accessible Chromatin (ATAC-seq, ^51^) on nuclei isolated separately from these populations of primary bovine articular chondrocytes. The 907 sites selectively accessible in SZCs were enriched for 4 distinct sequence motifs, including CRE (cAMP response elements; TGACGTCA) and TRE (TPA response elements; TGAGTCA) (Fig. 6A). Both these motifs bind Creb5, either as a homodimer or as a heterodimer with Jun ^40^. In contrast, ATAC-Seq peaks that were selectively accessible in DZCs were most highly enriched for 2 distinct sequence motifs, which serve as binding sites for either Tead or Runx transcription factors (Fig. 6B). Notably, among genes enriched in either SZCs or DZCs which contained transcription start sites located ≤25 kb from differentially accessible ATAC sites, we observed a high correlation between chromatin access and zone-specific gene expression (Fig. 6C). This correlation was highest at the *Prg4* locus (Fig. 6C), where 4 distinct regions (E1-E4) were selectively accessible in SZCs (Fig. 7A). Chromatin IP (ChIP) of iCreb5-HA DZCs with HA antibody, followed by PCR analysis of the precipitated DNA, revealed specific occupancy of Creb5 at the two *Prg4* promoter-proximal sites, E1 and E2, and increased binding induced by TGF-β treatment at both candidate *cis*-elements (Fig. 7B). Together, these observations implicate CRE/TRE binding TFs such as Creb5 in distinguishing the two articular chondrocyte populations. Atf2, which is closely related to Creb5, has been found to directly interact with Smad3/4 ^52^. We observed that Creb5 can similarly co-immunoprecipitate Smad2/3 (Supplementary Figure 3), suggesting that TGF-β signaling may in part increase *Prg4* expression by inducing nuclear translocation of Smad2/3 and consequent interaction with Creb5.

**Fig. 6.**
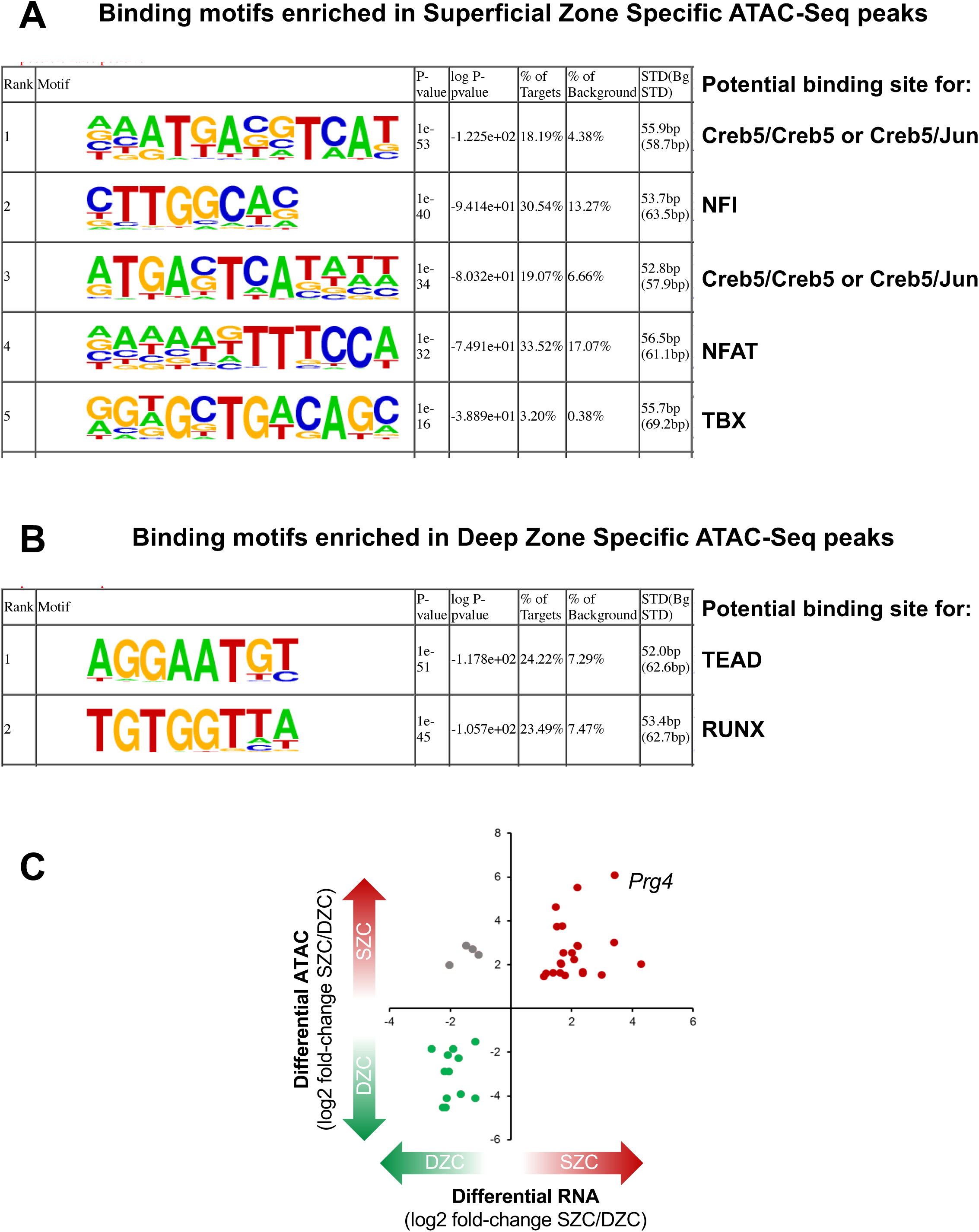
Creb5 binding motifs are enriched in SZC-specific ATAC-Seq peaks. (A) Homer motif analysis of ATAC-Seq peaks that are enriched in SZCs versus DZCs indicated that binding sites for Creb5, Nfi, Nfat, and Tbx TFs are enriched in superficial zone-specific ATAC-Seq peaks. (B) Homer motif analysis of ATAC-Seq peaks that are enriched in DZCs versus SZCs indicated that the binding sites for Tead and Runx TFs are enriched in deep zone-specific ATAC-Seq peaks. The frequency of these sequence motifs in zone-specific ATAC-Seq peaks (% of targets) versus their frequency in the genome (% background) is displayed. (C) A high correlation was observed between zone-specific ATAC-Seq sites (located ≤25 kb from the transcription start site) and zone-specific gene expression in both SZCs and DZCs. This correlation was highest at the *Prg4* locus, where 4 distinct regions (E1-E4) were selectively accessible in SZCs.

**Fig. 7.**
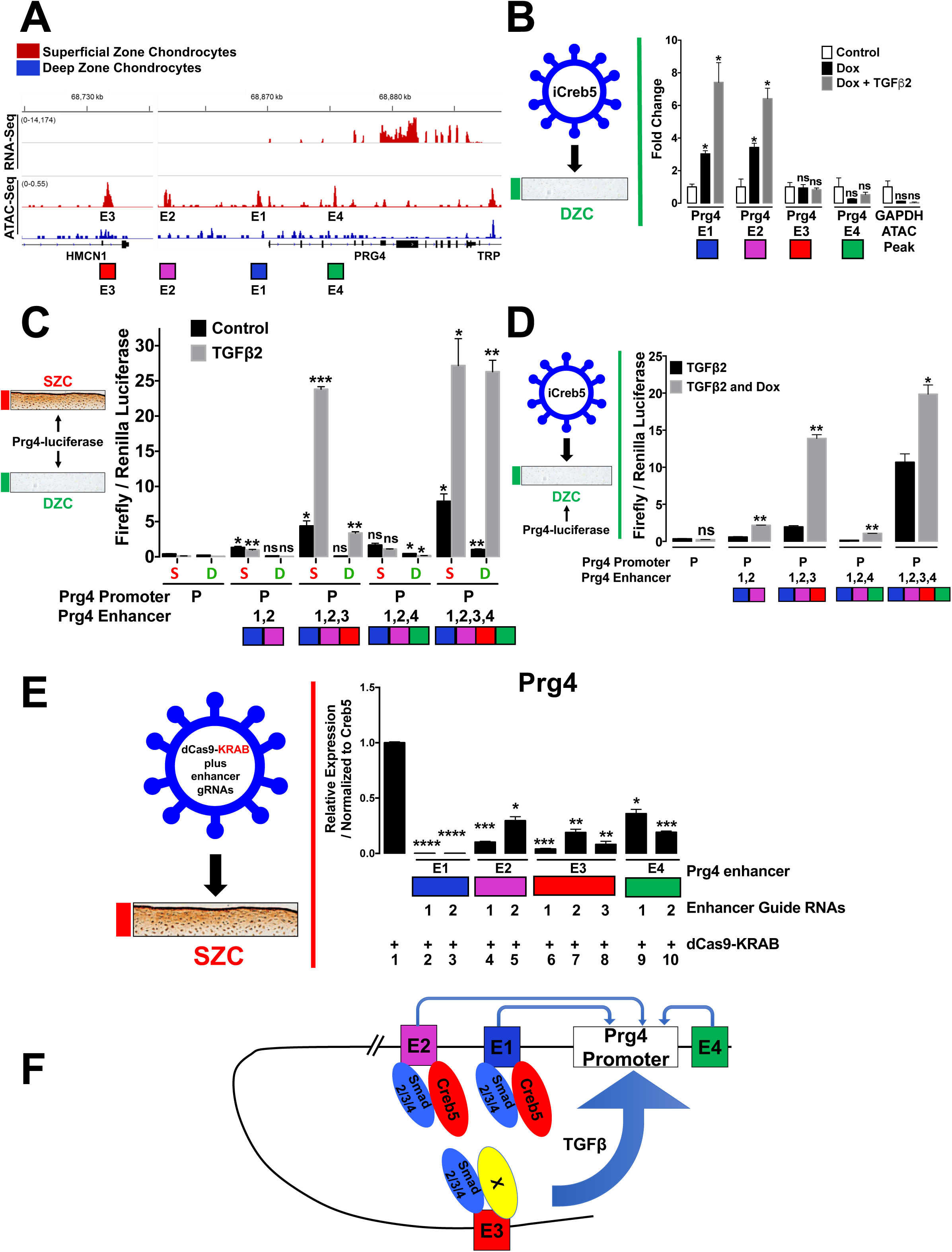
*Prg4* expression is regulated by the combinatorial activity of several regulatory elements. (A) Comparison of RNA-Seq (top tracks) and ATAC-Seq (bottom tracks) surrounding the *Prg4* locus. Signals for either SZCs (red) or DZCs (blue) are displayed. Putative *Prg4* regulatory elements (E1-E4) are indicated. (B) Bovine articular chondrocytes were infected with lentivirus encoding doxycycline-inducible HA-tagged Creb5 (iCreb5-HA). The cells were cultured in either the absence or presence of doxycycline and TGF-β2, as indicated. ChIP-qPCR was performed with anti-HA. Creb5-HA occupancy on ATAC-Seq peaks E1-E4 in the *Prg4* locus or in the GAPDH locus are displayed. (C) SZC-enriched ATAC-Seq peaks surrounding the *Prg4* locus work in combination. Either superficial zone (S) or deep zone (D) bovine articular chondrocytes were co-transfected with a firefly luciferase reporter driven by both the *Prg4* promoter plus a combination of enhancers (E1, E2, E3, and E4) surrounding this gene and a CMV-renilla luciferase construct. Relative expression of firefly/renilla luciferase is displayed (E1E2-*Prg4*-promoter construct compared to *Prg4*-promoter-luciferase; all rest compared to E1E2-*Prg4*-promoter luciferase). Similar results have been obtained in 3 independent biological repeats. (D) DZCs were infected with a lentivirus encoding doxycycline-inducible Creb5-HA. After selection in G418, the cells (iCreb5-DZCs) were transfected with Prg4-firefly luciferase expression constructs as described above and cultured with TGF-β2 (20 ng/ml) in either the absence or presence of doxycycline (1μg/ml), as indicated for 2 days. Relative expression of firefly/renilla luciferase is displayed. Similar results have been obtained in 2 independent biological repeats. (E) SZCs were infected with lentivirus encoding only dCas9-KRAB or with lentivirus programmed to encode both dCas9-KRAB plus guide RNAs targeting the various SZC-enriched ATAC-seq peaks (2-3 different guides for each ATAC-Seq peak) that surround the *Prg4* locus. After selection in puromycin, the cells were cultured in the presence of TGF-β2 (20 ng/ml) for 3 days. Gene expression was assayed by RT-qPCR and normalized to *Creb5*. Relative levels of *Prg4* expression in cells expressing dCas9-KRAB plus various guide RNAs (lanes 2-10) are compared to cells expressing only dCas9-KRAB (lane 1). Similar results have been obtained in 3 independent biological repeats. (F) Model for Creb5-dependent induction of *Prg4* expression.

As *Prg4* is a seminal SZC-specific marker, we examined whether SZC-specific sites E1-E4 function as enhancers. To this end, we appended each sequence upstream of the *Prg4* promoter and firefly luciferase cDNA; and assessed the constructs’ ability to drive reporter gene expression in primary bovine SZCs or DZCs. While the *Prg4* promoter alone showed little activity, addition of both E1 and E2 upstream elements drove luciferase expression specifically in SZCs, but TGF-β2 did not add to this effect (Fig. 7C). Thus, E1 and E2 both interact with Creb5-HA (in iCreb5-DZCs) and drive SZC-specific gene activity, but do not respond to TGF-β. In striking contrast, addition of the distal regulatory element E3 to the promoter-proximal E1 and E2 elements robustly induced TGF-β2 responsive luciferase expression, in both SZCs and DZCs (Fig. 7C). The E4 element alone failed to confer a TGF-β response and did not interfere with the potent activity of E3 (Fig. 7C). Lastly, while treatment of iCreb5-DZCs with doxycycline plus TGF-β2 weakly induced expression of the E2E1-Prg4-luciferase construct, the addition of the E3 element to this reporter strongly enhanced doxycycline-dependent luciferase activity (Fig. 7D). Thus, the E3 element, located 140 kb upstream of the *Prg4* transcription start site, contains an enhancer that responds robustly to TGF-β2 in both SZ and DZ chondrocytes.

To ask whether the SZC-enriched sites E1-E4 are necessary to drive *Prg4* expression in SZCs, we infected these cells with a lentivirus encoding either only a dead-Cas9-KRAB fusion protein (dCas9-KRAB) or dCas9-KRAB plus guide RNAs targeting each *cis*-element. Targeting of dCas9-KRAB to putative enhancers recruits the H3K9 methyltransferase SETDB1, hence increasing local H3K9me3 levels and repressing target genes ^53^. Compared to dCas9-KRAB alone, inclusion of guide RNAs against E1, E2, E3 or E4 significantly repressed TGF-β2 induction of *Prg4* (Fig. 7E). Notably, targeting of E1 with 2 different gRNAs repressed *Prg4* induction to 0.1% of control levels, while targeting of E2, E3, or E4 repressed *Prg4* to 4-36% of control levels, (Fig. 7E). While E1, E2 and E3 all lie upstream of the *Prg4* coding reagion, E4 is located in an intron of the *Prg4* gene. Thus, it is possible that recruitment of dCas9-KRAB to E4 and consequent deposition of H3K9me3 modification to the *Prg4* gene proper may block progression of RNA polymerase II, and thereby decrease production of *Prg4* mRNA. Taken together, these findings reveal that expression of *Prg4* in SZCs is dependent upon both the proximal *cis*-elements E1 and E2, which bind Creb5, and a distant E3 element, that drives TGF-β responsive gene activity.

## Discussion

In this work, we document that Creb5 is specifically expressed in the superficial zone of the articular cartilage and is necessary for TGF-β signaling to promote *Prg4* expression in superficial zone bovine articular chondrocytes. In developing mouse knees, Creb5 is expressed in the articular perichondrium, in superficial cells of the prospective meniscus, and in synovial fibroblasts that line the joint cavity; the same tissues that express *Prg4* ^2, 4, 44^. Thus, Creb5 is present in the tissues that express *Prg4* in murine, bovine and human joints, and is necessary to maintain competence for *Prg4* expression in superficial zone articular chondrocytes. In addition, when misexpressed in deep zone bovine articular chondrocytes, Creb5 confers the competence for TGF-β and EGFR signals to induce *Prg4* expression in these cells. The regionalized expression of Creb5 in the articular cartilage, which is confined to the superficial zone, helps to explain how *Prg4* expression is similarly constrained to this region of the articular cartilage. As nuclear-localized CREB5 is specifically present in the lubricin-expressing superficial zone of articular cartilage in adult human femoral head articular cartilage, it is plausible that this transcription factor may also be necessary to sustain robust lubricin expression in adult human articular cartilage. Loss of Creb5 function in SZCs decreased the expression of both *Prg4* and other SZC-specific genes (such as *Epha3*). Thus, it seems likely that *Creb5* may play a larger role in maintaining the unique biological properties of the superficial zone of the articular cartilage.

Prior work has indicated that mechanical motion can promote the expression of *Prg4* in articular cartilage via multiple *Creb1*-dependent, fluid flow shear stress-induced signaling pathways ^33^. In contrast to *Creb5*, whose expression is restricted to the superficial zone of the articular cartilage, *Creb1* transcripts are equally expressed in both superficial and deep zones of this tissue; but at a significantly lower level than *Creb5*. While both *Creb1* and *Creb5* can bind to overlapping binding sites ^40, 54^, their transcriptional activities are modulated by distinct signaling pathways. PKA-mediated phosphorylation of Creb1 (on Ser133) promotes the interaction of this TF with the KIX domain of the co-activator proteins CBP (CREB-binding protein) and/or p300 ^55–57^. Both PGE2 and PTHrP signaling pathways (or forskolin treatment), which can all activate PKA, promote *Prg4* expression in cultures of epiphyseal chondrocytes taken from 5-day-old mice ^33^. Despite its name, *Creb5* is most closely related to the ATF2/7 family ^40^, whose transcriptional activity is regulated by SAPK and MAPK signaling pathways (reviewed in ^42, 43^). Interestingly, the PKA activator 8-Br-cAMP can induce phosphorylation of ATF2 on T71 ^58^. As this phosphorylation site is conserved in Creb5, it is possible that PKA signaling may similarly be able to regulate Creb5 activity. Future ChIP-Seq analysis will be necessary to evaluate whether Creb1 and Creb5 share overlapping binding sites on *Prg4* regulatory elements, and whether these TFs work either synergistically or in parallel to activate the expression of *Prg4* in response to differing signaling pathways.

By performing both ATAC-Seq and chromatin-IP (ChIP) for Creb5, we have found that Creb5 directly binds to two *Prg4* promoter-proximal regulatory elements (E1 and E2), which display an open chromatin conformation specifically in superficial zone articular chondrocytes. Interestingly, the *Prg4* promoter-proximal regulatory elements (E1 and E2) which interact with Creb5, can drive superficial zone-specific chondrocyte gene expression, but cannot respond to TGF-β signals. In striking contrast, appending a distal 5’ regulatory element (E3), which also displays increased chromatin accessibility in superficial zone articular chondrocytes (but does not directly bind to Creb5), immediately adjacent to the more proximal Creb5 binding elements (in a luciferase reporter construct) drove robust luciferase expression in response to TGF-β2. Recruitment of a dead-Cas9-KRAB fusion protein to either the Prg4 promoter-proximal regulatory elements (that bind to Creb5) or to more distal regulatory elements, significantly blunted induction of *Prg4* by TGF-β signals in superficial zone chondrocytes. Our working hypothesis is that direct interaction of Creb5 with either E1 or E2 alters the chromatin structure of these regulatory elements, such that they can interact with more distal regulatory elements, which in turn drive robust TGF-*β−*dependent induction of *Prg4* (Fig. 7F). It may be relevant in this regard that Creb5 can bind to DNA as a heterodimer with Jun; and that Jun/Fos (in an AP1 complex) can recruit the SWI/SNF (BAF) chromatin remodeling complex to establish accessible chromatin on targeted enhancer elements ^59^. Future studies will be necessary to investigate whether Creb5 can similarly remodel chromatin to establish enhancer accessibility for the interaction of other transcription factors.

Consistent with the restricted expression of Creb5 in the superficial zone of the articular cartilage, we observed that Creb5 binding sites ^40^, including CRE (cAMP response elements; TGACGTCA) and TRE (TPA response elements; TGAGTCA), were both enriched in SZC-specific ATAC-Seq peaks. In contrast, binding sites for either Tead or Runx transcription factors were enriched in DZC-specific ATAC-Seq peaks. While it is currently unclear what regulates either the SZC-specific expression of *Creb5* or the occupancy of Tead and Runx transcription factors on DZC-specific accessible regions of chromatin, these findings underscore the differential transcriptional regulation in these two regions of the articular cartilage. Notably, *Prg4* expressing cells in embryonic and neonatal mice have been found to give rise to all regions of the articular cartilage in adult animals, including those in the deep zone of this tissue ^8–10^. Thus, the absence of *Creb5* expression in deep zone articular chondrocytes suggests that *Creb5*, which is initially expressed in the precursors of all articular chondrocytes, is somehow down-regulated together with *Prg4*/lubricin in the deep zone of the articular cartilage.

TGF-β ^36, 37^, EGFR ^38^, and Wnt/β-catenin ^11, 34, 35^ signaling have all been shown to be necessary to maintain expression of *Prg4* in the articular cartilage. Interestingly, we observed that TGF-β and EGFR signaling pathways augment the ability of either endogenous Creb5 or exogenous iCreb5 to promote expression of *Prg4* in either superficial or deep zone bovine articular chondrocytes, respectively. Binding of Creb5 to *Prg4* promoter-proximal regulatory elements (E1 and E2) was considerably enhanced by TGF-β administration, consistent with our finding that Creb5, like ATF2 ^52^, can be co-immunoprecipitated with Smad2/3. Thus, TGF-β signaling may increase occupancy of Creb5 on the two *Prg4* promoter-proximal enhancer elements (E1 and E2) by inducing nuclear translocation of Smad2/3 and consequent interaction with Creb5. Future studies will be necessary to determine whether the requirement for Wnt ^11, 34, 35^ and EGFR ^38^ signals to maintain the expression of *Prg4* in articular cartilage is due to modulation of either the expression or activity of Creb5, or of other necessary co-factors. Exogenous Creb5 could drive relatively high-level expression of *Prg4* (approximately equal to 75% of *Gapdh* transcript levels) in deep zone bovine articular chondrocytes treated with TGF-β. In contrast, the combination of *Creb5* and TGF-β signaling only induced *PRG4* expression to approximately 0.2% of *GAPDH* transcript levels in either a human chondrosarcoma cell line (SW1353) or in an immortalized human costal chondrocyte cell line (C-28/12)^49^. Thus, it seems likely that in addition to Creb5, other transcriptional regulators that are unique to articular chondrocytes may play a role in driving high-level level expression of *Prg4.* In addition to Creb5, we found that 28 transcriptional regulators were more highly expressed (at least 2-fold) in SZCs than in DCZs; and only one TF (Sall1) was more highly expressed in DZCs (Table 1). Notably, Nfat binding motifs were significantly enriched in SZC-specific ATAC-Seq peaks, suggesting that members of the Nfat family may cooperate with Creb5 to promote SZC-specific gene expression. Indeed, cartilage-specific deletion of *Nfatc1 in Nfatc2*-null mice leads to early onset osteoarthritis, and decreased expression of *Prg4* ^32^. Future studies will be necessary to determine whether either Nfat TFs or any of the other SZC-enriched transcriptional regulators play a direct role in driving the expression of either *Prg4* or other SZC-specific genes.

Creb5 shares a high degree of homology with both Atf2 and Atf7 in both its DNA binding domain and its N-terminus, which contains two highly conserved Threonine-Proline sequences (T59 and T61 in Creb5) that are substrates for p38 kinase, JNK, and ERK (reviewed in ^42, 43^). We have found that phosphorylation of Creb5(T61) can be boosted by both EGFR and TGF*−β* signals in superficial zone chondrocytes and that phosphorylation of Creb5(T61) is blocked in TGF-β treated iCreb5-DZCs by either a p38 inhibitor (SB203580) or a JNK inhibitor (SP600125). In addition, we noted that Stress Activated Protein Kinase (SAPK) function is required to promote *Prg4* expression in either superficial zone chondrocytes (that express endogenous Creb5) or iCreb5-DZCs (programmed to express iCreb5). Substitution of alanine for threonine in the two highly conserved SAPK phosphorylation sites of either Atf2 or Atf7 cripples the activity of the adjacent N-terminal transcriptional activation domain of these proteins ^60, 61^ and the biological activity of Atf2 in vivo ^62^. In striking contrast, we found that similar mutations in the SAPK kinase sites of Creb5 (i.e., iCreb5-T59,61A) did not depress the ability of this transcription factor to induce *Prg4* expression in deep zone chondrocytes treated with either an EGFR ligand and/or TGF-β signals. These findings indicate that while Stress Activated Protein Kinases can promote phosphorylation of Creb5 (on T59, T61), induction of *Prg4* expression, by EGFR and TGF-β signals, is dependent upon SAPK-mediated phosphorylation of either other sites in Creb5 or phosphorylation of other substrates. The transcriptional activity of a chimeric protein containing the GAL4 DNA binding domain fused to the N-terminus of Creb5 (GAL4-Creb5-(1-128) was attenuated by simultaneous T59A/T61A mutations. Thus, it will be interesting to determine whether, in contrast to *Prg4*, the induction of other Creb5 target genes are regulated by phosphorylation of T59 and T61 in Creb5.

## Methods

### Isolation and culture of bovine articular chondrocytes

The knee joints from 1-2 week old bovine calves were obtained from Research 87 (a local abattoir in Boylston, MA) directly after slaughter. The intact femoropatellar joints was isolated by transecting the femur and mounting the distal segment in a drilling apparatus. The femoropatellar articular cartilage was then exposed by opening the joint capsule, severing the medial, lateral, and cruciate ligaments, and removing the tibia, patella, and surrounding tissue. Four to six cylindrical cores of cartilage and underlying bone, 9.5 mm in diameter and −15 mm deep, were drilled from each facet (medial and lateral) of the femoropatellar groove. During this entire process, the cartilage was kept moist and free of blood by frequent rinsing with sterile PBS supplemented with antibiotics (100 U/ml penicillin and 100 μg/ml streptomycin). Each core was then inserted into a cylindrical sample holder for a sledge microtome (Model 860, American Optical, Buffalo, NY). An initial ∼200-300 micron thick slice of superficial zone articular cartilage (SZC) was first harvested via the microtome. The next approximately 4 mm thick slice of middle zone cartilage tissue was then removed from the same core and discarded. Finally, an 800 micron to 1 mm thick slice of deep zone articular cartilage (DZC) was then harvested from the same core. All of the superficial zone slices from a given knee joint were placed in a 50 mL centrifuge tube filled with medium (DMEM with 10 mM HEPES, 0.1 mM nonessential amino acids, and additional 0.4 mM proline, 25 pg/ml ascorbate) and supplemented with 10% fetal bovine serum. Similarly, all deep zone slices from a given joint were placed in a separate tube filled with medium and serum. To isolate chondrocytes from the superficial and deep zone slices, deep zone cartilage shavings were chopped into pieces about 1 mm^3^. There was no need to chop superficial zone shavings as they are thin enough. Both superficial zone and chopped deep zone cartilage were digested in 10 ml of pronase (1 mg/ml) in DMEM with 1% penicillin & streptomycin for 1 hour. Pronase was replaced by collagenase D (1 mg/ml) in DMEM with 1% penicillin & streptomycin and cartilage tissue was digested in the incubator at 37 degrees overnight. The next day, the dissociated cells were filtered through a 70 μm strainer, counted, pelleted for 5 min at 1200rpm and resuspended in DMEM/F12 plus 10% FBS. For lentivirus infection the cells were plated into a 6-well plate (2-3 million cells per well). 24 hours after plating the medium was changed with new DMEM/F12 plus 10% FBS.

### RNA-Seq Analysis

Newly isolated bovine superficial zone chondrocytes and deep zone chondrocytes were cultured (in DMEM/F12 plus 10% FBS) for three days prior to performing RNA-Seq analysis. RNA from superficial zone chondrocytes and deep zone chondrocytes were purified using Trizol reagent (Life Technologies). Genomic DNA in RNA samples was removed using TURBO DNA-free™ Kit (Thermo Fisher Scientific, Cat#: AM1907). Total RNA (5 to 10 ng) was purified using the manufacturer’s instructions and used to prepare libraries with SMART-Seq v4 Ultra Low Input RNA Kit (Clontech) followed by sequencing on a NextSeq 500 instrument (Illumina) to obtain 75-bp single-end reads. Raw RNA-seq reads were assessed with FastQC 0.11.3 followed by MultiQC 1.2 aggregation ^63^ to determine sequence quality, per-base sequence quality, per-read GC content (∼50), and per base N content. Read pairs were aligned to the Bos taurus genome Ensembl build UMD 3.1 version 88 using STAR 2.5.2b ^64^ employing a custom index, with read counting for an unstranded library preparation. Counts were normalized using Trimmed Means of M-values (TMM) ^65^ as part of the edgeR package ^66, 67^, and modelled for biological and gene-wise variation. Differential expression between sample types was determined through the Exact Test in edgeR, with Benjamini-Hochberg ^68^ multiple testing correction (FDR) set at < 0.05.

### ATAC-Seq Analysis

Newly isolated bovine superficial zone articular chondrocytes and deep zone articular chondrocytes were cultured (in DMEM/F12 plus 10% FBS) for three days prior to performing ATAC-Seq analysis. ATAC-Seq ^51, 69^ was performed on replicate samples of 8,000 to 35,000 superficial zone chondrocytes or deep zone chondrocytes. Cultured chondrocytes were first digested into single cells using trypsin, and then trypsin was neutralized by serum. Digested single cells were washed twice in ice-cold PBS, resuspended in 50 μl ice-cold ATAC Lysis Buffer (10 mM Tris·Cl, pH 7.4, 10mM NaCl, 3mM MgCl_2_, 0.1% (V/V) Igepal CA-630), and centrifuged at 500 g at 4 °C to isolate nuclear pellets. Nuclear pellets were treated with Nextera Tn5 Transposase (Illumina, FC-121-1030) in a 50 μl reaction for 30 min at 37 °C. Transposed DNA was immediately isolated using a Qiagen MinElute PCR Purification Kit, and then PCR amplified in a 50 μl reaction using a common forward primer and different reverse primers with unique barcodes for each sample as per ^69^. After 5 cycles of PCR, 45 μl of the reaction was kept on ice; while 5 μl reaction was amplified by RT-qPCR for 20 cycles to determine the cycles required to achieve 1/3 of the maximal RT-qPCR fluorescence intensity. The remaining 45 μl of the reaction was then amplified by 8 additional cycles to achieve 1/3 of the maximal RT-qPCR fluorescence intensity (as determined above). The amplified DNA was purified using a Qiagen MinElute PCR Purification Kit and primer dimers (<100 bp) were removed using AMPure beads (Beckman Coulter). ATAC-Seq was performed with two biological repeats to ensure the robustness of the data sets. Raw ATAC-Seq reads were aligned to the bovine genome (Bostau 6) using Bowtie2 ^70^. Aligned signals in raw (bam) files were filtered to remove PCR duplicates and reads that aligned to multiple locations. Peaks were identified using MACS v1.4 ^71^.

### Construction of lentivirus encoding shRNA targeting Creb5

Bovine Creb5 shRNAs targeting the Creb5 3’UTR were designed using Block-iT RNAi designer (Thermo Fisher). The most efficient Creb5 shRNA sequence we identified was: CCG G<u>GC CTT CAA GAA GAG CTG TTG C</u>CT CGA G<u>GC AAC AGC TCT TCT TGA AGG C</u>TT TTT G (targeting sequence is underlined; employed in Fig. 3A-C). Oligos (ordered from Integrated DNA Technologies) were annealed in a 10 μl reaction (1 μl Forward oligo (100 μM), 1μl Reverse oligo (100 μM), 1μl T4 ligation buffer (10X), 6.5 μl nuclease-free H_2_O, 0.5 μl T4 Polynucleotide Kinase (NEB)) in a PCR machine, programmed to cycle: 37 °C 30 min, 95 °C 5 min, and then ramp down to 25 °C at 5 °C/min. Annealed oligos that are compatible with the sticky ends of EcoRI and AgeI, were diluted (1:100) and cloned into pLKO.1 TRC-Cloning vector (Addgene # 10878) that had been digested with EcoRI and AgeI, and gel purified. Lentiviral vector encoding shRNA targeting bovine Creb5 (shCreb5) was verified by sequencing (Genewiz). Lentiviral control vector containing a scrambled shRNA (shSCR) was ordered from Addgene (scramble shRNA, Addgene # 1864).

### CRISPR/Cas9 targeting the DNA binding domain of *Creb5*

The CRISPR/Cas9 system was used to introduce insertions /deletions (indels) into the *Creb5* DNA binding domain in bovine superficial zone chondrocytes. Briefly, sequence specific sgRNAs that guide Cas9 to the genomic region encoding the *Creb5* DNA binding domain were designed following the instructions located at (http://crispr.mit.edu/). The most efficient guide targeting the DNA binding domain of bovine *Creb5* that we identified is: C TGA AGC TGC ATG TTT GTC T (employed in Fig. 3D-F). Oligos (ordered from Integrated DNA Technologies) were annealed in a 10 μl reaction (1 μl Forward oligo (100 μM), 1μl Reverse oligo (100 μM), 1μl T4 ligation buffer (10X), 6.5 μl nuclear-free H_2_O, 0.5 μl T4 Polynucleotide Kinase (NEB)) in a PCR machine programmed to cycle: 37 °C 30 min, 95 °C 5 min, and then ramp down to 25 °C at 5 °C/min. The annealed oligos were diluted (1:200) and cloned into lentiCRISPRv2 (Addgene # 52961) using a Golden Gate Assembly strategy (containing: 100 ng circular lentiCRISPRv2, 1 μl diluted oligo, 0.5 μl BsmBI (Thermo Fisher), 1 μl Tango buffer (10X), 0.5 μl DTT (10mM), 0.5 μl ATP (10 mM), 0.5 μl T4 DNA Ligase (NEB), H_2_O up to 10 μl) with 20 cycles of: 37 °C 5 min, 21 °C 5 min. The ligation reaction was then treated with PlasmidSafe (Epicentre, Cat#: E3101K) to digest any residual linearized DNA. PlasmidSafe treated plasmid was transformed into Stbl3 competent cells (Thermo Fisher). LentiCRISPRCreb5gRNA plasmid containing the Creb5 gRNA was verified by sequencing (Genewiz). The parental vector, lentiCRISPRv2, was used to generate control lentivirus.

### T7 endonuclease I (T7E1) assay to detect indels in *Creb5*

The T7 endonuclease I (T7E1) assay was used to detect on-target CRISPR/Cas9 induced insertions and deletions (indels) in cultured cells. cDNA derived from mRNA isolated from either lentiCRISPRv2 or lentiCRISPRCreb5gRNA infected bovine superficial zone chondrocytes was employed for the T7E1 assay. A 210bp cDNA fragment, which flanks the Creb5 gRNA cleavage site, was amplified using Creb5 RT-qPCR primers (Supplementary Table 2) in a 25 μl PCR reaction (12.5 μl Q5 High-Fidelity 2X Master Mix, 1.25 μl Forward Primer, 1.25 μl Reverse Primer, cDNA, nuclease-free H_2_O to a final volume of 25 μl). After denaturation of the cDNA at 98 °C 30s; the PCR machine was programed to cycle 35 times at: 98 °C 10 s, 60 °C 15 s, 72 °C 15 s; 72 °C 2 min; 4 °C hold. The PCR product was denatured and annealed in an 18 μl reaction (15 μl PCR product, 2 μl NEB Buffer2 (10X), 1 μl nuclease-free H_2_O) with the following the cycling conditions: 95 °C 10 min; 95-85 °C (ramp rate of - 2°C/sec); 85-25 °C (ramp rate of - 2°C/sec). After denaturing and reannealing the PCR products, T7 endonuclease I (2 μl) was added, and the mixture was incubated at 37 °C for 60 min. Cleavage of the PCR products by T7 endonuclease was assayed by agarose gel electrophoresis.

### Generation of lentivirus encoding either WT or mutant iCreb5

Total RNA (containing *Creb5* transcripts) was isolated from bovine superficial zone articular chondrocytes using Trizol reagent (Life Technologies). cDNA was reverse transcribed using the oligo dT reverse transcription kit SuperScript® III First-Strand Synthesis System (Life Technologies, cat. no. 18080051). The bovine *Creb5* open reading frame (508aa; see NM_001319882.1) was predicted by RNA-Seq of bovine superficial zone articular chondrocytes. A 1524 bp cDNA fragment (encoding 508aa) was PCR amplified from bovine superficial zone articular chondrocyte cDNA using the Q5 High-Fidelity 2X Master Mix (NEB, cat. no. M0492S) and then cloned into the pCR®-Blunt vector (Life Technologies, Cat#: K270020). Using the pCR®-Blunt-bovine Creb5 as a template, a fragment of DNA encoding 3xHA tags was added onto the C-terminus of Creb5 (immediately before the stop codon). HA-tagged Creb5 was then cloned into a Gateway vector (pENTR-Creb5-HA) using the pENTR™/SD/D-TOPO® Cloning Kit (Life Technologies, Cat#: K2420-20). Creb5-HA was then transferred from pENTR-Creb5-HA into the pInducer20 (Addgene # 44012) lentivirus destination vector or into the pLenti CMV Puro DEST (w118-1) vector (Addgene Plasmid #17452) using Gateway LR Clonase (Thermo Fisher, cat. no. 11791020), to generate pInducer20-iCreb-WT or pLenti-Creb5, respectively. pLenti-GFP vector (EX-EGFP-Lv102) was purchased from GeneCopoeia. To generate lentivirus vectors encoding iCreb5 mutants, the DNA sequence encoding T59/T61 sites in pENTR-Creb5-HA was mutated into sequence encoding either T59/T61A or into T59/T61D, respectively, using the Q5® Site-Directed Mutagenesis Kit (NEB, Cat#: E0554S) following the manufacturer’s instructions. Creb5 mutants were then cloned into the lentivirus destination vector pInducer20 using Gateway technology.

### Lentivirus encoding dCas9-KRAB targeting *Prg4* ATAC-seq peaks

The CRISPR interference (CRISPRi) system was used to study the function of *Prg4* enhancer elements (E1, 2, 3, 4) in bovine superficial zone chondrocytes. Briefly, sequence specific sgRNAs that direct a dead-Cas9-KRAB fusion protein (dCas9-KRAB; ^53^) to the genomic region of the *Prg4* enhancer elements were designed following the instructions at CHOPCHOP (http://chopchop.cbu.uib.no/). Oligos (ordered from Integrated DNA Technologies) were annealed in a 10 μl reaction (1 μl Forward oligo (100 μM), 1μl Reverse oligo (100 μM), 1μl T4 ligation buffer (10X), 6.5 μl nuclear-free H_2_O, 0.5 μl T4 Polynucleotide Kinase) in a PCR machine programmed to cycle: 37 °C 30 min, 95 °C 5 min and then ramp down to 25 °C at 5 °C/min. The annealed oligos were diluted (1:200) and cloned into pLV hU6-sgRNA hUbC-dCas9-KRAB-T2a-Puro (encoding dCas9-KRAB; Addgene # 71236) using a Golden Gate Assembly strategy (containing: 250 ng circular pLV hU6-sgRNA hUbC-dCas9-KRAB-T2a-Puro, 1 μl diluted oligo, 0.5 μl BsmBI (Thermo Fisher), 1 μl Tango buffer (10X), 0.5 μl DTT (10mM), 0.5 μl ATP (10 mM), 0.5 μl T4 DNA Ligase (NEB), H_2_O up to 10 μl) with 20 cycles of: 37 °C 5 min, 21 °C 5 min. The ligation reaction above was then treated with PlasmidSafe (Epicentre, Cat#: E3101K) to digest any residual linearized DNA. PlasmidSafe treated plasmid was transformed into Stbl3 (Thermo Fisher) competent cells. dCas9-KRAB plasmids containing the various Prg4 enhancer gRNAs were verified by sequencing (Genewiz). The gRNAs targeting the bovine Prg4 enhancer elements are listed in Supplementary Table 5.

### Growth and purification of lentivirus

HEK293 cells were used to package lentivirus. HEK293 cells were plated in a 15-cm dish in 25ml of DMEM/F12 (Invitrogen) supplemented with 10% heat-inactivated fetal bovine serum (Invitrogen) and Pen/Strep at 37°C with 5% CO_2_. Transfection was performed when the cells were approximately 70–80% confluent. Lentiviral expression plasmid (6 μg), psPAX2 (4.5 μg, Addgene #12260), and pMD2.G VSVG (1.5μg, Addgene #12259) plasmids were added into a sterile tube containing 500 μl of Opti-MEM® I (Invitrogen). In a separate tube, 36 μl of Fugene 6 was diluted into 500 μl of Opti-MEM I. The diluted Fugene 6 reagent was added drop-wise to the tube containing the DNA solution. The mixture was incubated for 15–25 minutes at room temperature to allow the DNA-Fugene 6 complex to form. The DNA-Fugene 6 complex was directly added to each tissue culture dish of HEK293 cells. After cells were cultured in a CO2 incubator at 37°C for 12-24 hours, the medium (containing the DNA-Fugene 6 complex) was replaced with 36 ml fresh DMEM/F12 medium supplemented with 10% heat-inactivated fetal bovine serum and penicillin-streptomycin. Cells were again placed in the CO2 incubator at 37°C; and virus-containing culture medium was collected in sterile capped tubes 48, 72 and 96 hours post-transfection. Cell debris was removed by centrifugation at 500 g for 10 minutes and filtration through nylon low protein-binding filters (SLHP033RS, Millipore). Virus was concentrated by ultracentrifugation (employing a SW32 rotor at 25K for 2hr and 30min at 4° C) and stored at −80° C.

### Infection of chondrocytes with lentivirus and RT-qPCR

Newly isolated bovine superficial zone articular chondrocytes, deep zone articular chondrocytes, SW1353 cells or immortalized human costal chondrocyte cells (C-28/I2; ^49^) were cultured for at least 2-3 days before infection. SW1353 cells were obtained from ATCC (HTB 94). The immortalized human costal chondrocyte cell line (C-28/I2)^49^ was obtained from Dr. Mary Goldring (Hospital for Special Surgery, Weill Cornell Medical College & Weill Cornell Graduate School of Medical Sciences). Ultracentifuge-concentrated lentivirus was added into DMEM/F12 medium (with 10% FBS) containing 7.5 μg/ml DEAE-Dextran (to increase infection efficiency). 24 hours after infection, medium was replaced with new DMEM/F12 medium (with 10% FBS) containing either 0.8 μg/ml puromycin or 500 μg/ml G418, for selection. After selection (for either 5 days in puromycin or 11 days in G418) the cells were re-plated onto low attachment tissue culture plates (Corning #3471 or #3473) in DMEM/F12 medium (with 10% FBS), without either puromycin or G418. After two to three days culture, RNA was harvested using Trizol reagent and cDNA were synthesized using SuperScript™ III First-Strand Synthesis SuperMix (Invitrogen, Cat#: 11752-050) according manufacture’s guidelines. RT-qPCR primers were synthesized by Integrated DNA Technologies. RT-qPCR was performed in an Applied Biosystem 7500 Fast Real-Time PCR Machine. Each experiment was performed with 2-3 biological repeats and each RT-qPCR assay was performed with technical repeats. Prism statistical software (GraphPad) was employed to analyze the data. Statistical analysis was performed using either unpaired or paired t tests. In RT-qPCR experiments, gene expression was normalized to that of either *GAPDH* or *Creb5*, as indicated. Gene expression was analyzed by RT-qPCR, employing at least one primer which was composed of sequences encoded by distinct exons. All RT-qPCR primers are listed in Supplementary Table 2 and Supplementary Table 11.

### Construction of GAL4-Creb5 fusion constructs

Using the pCR®-Blunt-bovine Creb5 cDNA as the template, Creb5 (1-128) and Creb5 (1-508) fragments were PCR amplified to lie between a 5’ EcoR1 and 3’ Xba1 site, using Q5 High-Fidelity 2X Master Mix (NEB, Cat#: M0492S). These amplicons were then cloned into EcoR1/Xba1 cleaved Gal4-HA-HIF1α (530-652) plasmid (Addgene # 24887), following the removal of the EcoRI-HA-HIF1α-XbaI fragment from this plasmid. The resulting plasmids were GAL4-Creb5(1-128) and GAL4-Creb5(1-508). GAL4-Creb5(1-128)T59/T61A and GAL4-Creb5(1-128)C18/C23S were generated by inducing point mutations in the GAL4-Creb5(1-128) vector using the Q5® Site-Directed Mutagenesis Kit (NEB, Cat#: E0554S). The control vector GAL4(1-147) was generated by introducing a stop codon immediately after the GAL4(1-147) sequence in the GAL4-Creb5(1-128) vector. All the plasmids were verified by sequencing.

### *Prg4*-luciferase constructs and luciferase assays

*Prg4* enhancer/promoter-firefly luciferase constructs were constructed by cloning various *Prg4* regulatory fragments into pGL4.10[*luc2*] vector (Promega). Bovine genomic DNA was extracted from bovine chondrocytes and was used as the template to amplify *Prg4* regulatory regions. A 543 bp fragment encoding the *Prg4* promoter and 5’ UTR up to the start codon of *Prg4* was amplified (with flanking BglII-HindIII sites) and cloned into pGL4.10[*luc2*] to generate pGL4.10 *Prg4*-Promoter-luciferase. A 1061 bp fragment encoding *Prg4* Enhancer1, the *Prg4* promoter and the 5’ UTR up to the start codon of *Prg4* was amplified (with flanking BglII-HindIII sites) and cloned into pGL4.10[*luc2*] to generate pGL4.10 E1-*Prg4* promoter-luciferase. A 585bp *Prg4* Enhancer2 fragment was amplified (with flanking EcoRV-BglII sites) and cloned into pGL4.10 E1-*Prg4* promoter-luciferase to generate pGL4.10 E2E1-*Prg4* promoter-luciferase. A 802 bp *Prg4* Enhancer3 fragment was amplified (with flanking XhoI-EcoRV sites) and cloned into pGL4.10 E2E1-*Prg4* promoter-luciferase to generate pGL4.10 E3E2E1-*Prg4* promoter-luciferase. A 331bp *Prg4* Enhancer4 fragment was amplified (with flanking NheI-XhoI sites) and cloned into pGL4.10 E3E2E1-*Prg4* promoter-luciferase to generate pGL4.10 E4E3E2E1-*Prg4* promoter-luciferase. High fidelity DNA polymerase (NEB, cat. no. M0492S) was used to generate all the constructs. All the plasmids were verified by sequencing. Prg4 Promoter and Enhancer 1, 2, 3, 4 sequences are listed in Supplementary Table 4.

*Prg4* enhancer/promoter-firefly luciferase reporters were co-transfected together with the pGL4.75 [hRlu/CMV] renilla luciferase reporter (Promega) into either superficial zone chondrocytes, deep zone chondrocytes, or deep zone chondrocytes previously infected with iCreb5WT lentivirus, using Lipofectamine ^TM^ LTX (Thermo Fisher, Cat#: 15338030). Twelve hours after transfection, cell culture medium was changed into new medium to remove the Lipofectamine DNA complex. After another twelve hours culture, transfected cells was re-plated into ultra low attachment tissue culture dishes (Corning, Cat#: 3471) for an additional 48 hours culture, in the either the absence or presence of TGF-β and Doxycycline. Cells were harvested and luciferase activity was measured using the Dual-Luciferase® Reporter Assay System (Promega, Cat#: E1960) in a Turner Biosystems Modulus Microplate Reader. Each experiment was performed with 2-3 biological repeats and each luciferase assay was performed with technical repeats. Prism statistical software (GraphPad) was employed to analyze the data. Statistical analysis was performed using either unpaired or paired t tests.

### Chromatin IP (ChIP)-PCR Analysis

Bovine deep zone articular chondrocytes infected with iCreb5-HA lentivirus were cultured in ultra low attachment tissue culture dishes (Corning, Cat#: 3471) for 2-3 days in DMEM/F12 supplemented with 10% FBS, in either the absence or presence of Doxycycline or Doxycycline plus TGF-β2. The chondrocytes were gently centrifuged (200g, 5 minutes), and digested with 0.25% Trypsin-EDTA (Thermo Fisher, Cat#: 25200056) at 37°C for seven minutes (until the tissue was dispersed into single cells). Trypsinization was stopped by adding 10x volume 10% FBS/DMEM.

Digested single cells were fixed in 1% Formaldehyde at room temperature for 20 min with shaking. Formaldehyde was quenched with 125 mM glycine on a shaking platform at room temperature for 10 min. Cells were then washed two times with cold PBS containing protease inhibitor cocktail (Roche) followed by lysis with 500 μl ChIP sonication buffer (0.5% SDS, 10mM EDTA, 50mM Tris-HCl pH 8.1). Chromatin (from ∼5 million cells in 500 μl ChIP sonication buffer) was sonicated with a Covaris E220 Sonication Machine (PIP=140, CBP=200, DF=5%, Avg Power =7.0, 300s each time, 6 times (Total 30min). Debris was removed by spinning samples 8 minutes at full speed at 4 °C. The chromatin was diluted with ChIP Dilution Buffer (1.1% TritonX-100, 1.2 mM EDTA, 16.7 mM Tris-HCL pH 8.1, 167 mM NaCl) up to 1500ul in 1.5ml eppendorf tube (DNA low bind tube). SDS concentration was adjusted to below 0.2% and TritonX-100 concentration was adjusted to 1%. Five micrograms of antibody (against the antigen of choice) was added to the chromatin and immunoprecipitated overnight at 4 °C. Protein A (15 μl) and Protein G (15 μl) beads were washed two times with ChIP Dilution Buffer and then added to the chromatin. The bead-chromatin complex was rotated at 4 °C for 2 hours and then washed two times each with Low Salt Buffer (0.1% SDS, 1% TritonX-100, 2mM EDTA, 20mM Tris-HCl pH 8.1, 150mM NaCl), High Salt Buffer (0.1% SDS, 1% TritonX-100, 2mM EDTA, 20mM Tris-HCl pH 8.1, 500mM NaCl), Lithium Buffer (0.25M LiCl, 1% IGEPAL CA630 (or NP-40), 1% Deoxycholic acid (Sodium Salt), 1mM EDTA, 10mM Tris pH8.1), and finally TE Buffer (10mM Tris-HCl pH8.0, 1mM EDTA). Chromatin-antibody complexes were eluted twice from beads using 100 μl of Elution Buffer (1% SDS, 0.1M NaHCO3) by incubation at room temperature for 10 minutes. NaCl was added into the 200 μl elute to a final 370 mM concentration. Cross-linked chromatin was reversed at 65 °C with shaking overnight followed by adding RNase A (to a final concentration of 0.2 mg/ml) and incubating at 37°C for one hour to digest RNA; and then adding Proteinase K (to a final concentration of 0.1 mg/ml) and incubating at 55 °C for one hour to digest protein. DNA fragments were purified using the PCR-purification kit with Min-Elute Columns (Qiagen) and analyzed using qPCR. In all ChIP-qPCR results, the level of immunoprecipitated DNA was normalized to that of the input DNA. Antibodies employed for ChIP are listed in Supplementary Table 7. Primers employed for ChIP-qPCR are listed in Supplementary Table 3.

### Co-Immunoprecipitation

Bovine deep zone articular chondrocytes were infected with a lentivirus encoding doxycycline-inducible Creb5 (3xHA tags was added onto the C-terminus of Creb5). After selection in G418, six million cells were cultured in low attachment tissue culture plates (Corning #3262) in DMEM/F12 medium (with 10% FBS, 20ng/ml TGF-β2) in the presence or absence of doxycycline for additional three days. Cells were harvested and suspended in 1.5 ml Mammalian Cell Lysis Buffer (MCLB, 50 mM Tris, pH 7.5, 150 mM NaCl, 0.5 % NP40,10 mM NaF) with Protease Inhibitors (PIs) Cocktail. The suspension was lysed by gently rocking at 4 °C for 1 hour and spun down at max speed on a table top centrifuge for 20 minutes at 4 °C. In 1.5 ml clear lysate, 100 μl lysate was kept as input and 25 μl of monoclonal anti-HA agarose was added into the remaining 1.4 ml sample. The binding was performed on a rocker at 4 °C overnight. After the binding, the agarose was washed with MLCB plus PIs for at least 5 times. The precipitated protein was eluted with NuPAGE LDS sample buffer (Invitrogen) plus NuPAGE Sample Reducing Agent (Invitrogen). The anti-HA agarose immunoprecipitated protein was probed with anti-Smad2/3 antibody by western blot. The input protein was probed with anti-Smad2/3 antibody and anti-HA antibody (Supplementary Table 6).

### Western Blots

Bovine superficial zone or deep zone articular chondrocytes were collected and lysed in Lysis Buffer (containing 50mM Tris. HCl, pH 7.4, 1% NP-40, 0.5% Sodium deoxycholate, 1% SDS, 150 mM NaCl, 2mM EDTA, 50mM NaF, protease inhibitor cocktail (Roche)). The cell lysate were incubated on ice for 15 min, and centrifuged at 3000g, at 4 °C to remove cell debris. Protein concentration was measured with a Bio-Rad Protein Assay Kit (BIO-RAD, Cat#: 500-0006). The protein was denatured at 70 °C for 10 min with LDS Sample Buffer (Invitrogen, Cat#: NP0007) and Sample Reducing Agent (Invitrogen, Cat#: NP0009). General SDS-PAGE processing procedures were followed. The blots were visualized by the enhanced chemiluminescence detection method, employing the Pierce ECL Western Blotting Substrate (Pierce, Cat#: 32106). All primary antibodies used for Western blots are listed in Supplementary Table 6.

### Immunohistochemical analysis

Mouse forelimb, hindlimb or human articular cartilage (procured by the National Disease Research Interchange (NDRI)) were dissected and fixed in 4% paraformaldehyde (in PBS) at 4°C overnight, washed with PBS, and incubated in 30% sucrose at at 4°C overnight. Tissues were embedded in OCT and frozen sections were cut at 12 μm using a cryostat. Any non-specific binding of primary antibodies was blocked by incubation with PBS containing 0.2% Tween-20 and 5% non-immune goat serum for 1 hour at room temperature. Primary antibody incubation was carried out at 4°C overnight with a rabbit polyclonal antibody to Creb5 (PA5-65593, Thermo Fisher, 1:100), a mouse monoclonal antibody to Prg4 (MABT401, EMD Millipore, 1:50), a rat monoclonal antibody to the HA epitope (11867423001, Sigma, 1:100), or a rabbit polyclonal antibody to Sox9 (AB5535, EMD Millipore, 1:100). After washing in PBST (3 × 15 minutes at room temperature), sections were incubated with Alexa Fluor 488 or 594 conjugated secondary antibodies (Thermo Fisher, 1:250) for 1 hour at room temperature. 4,6-diamidino-2-phenylindole (DAPI; 1μg/ml) was included with the secondary antibodies to stain DNA. Slides were washed in PBST (3 × 15 minutes at room temperature) and mounted under a coverslip with Aqua-mount (13800; Lerner Labs). Images were taken at the Nikon Imaging Center at Harvard Medical School. All antibodies employed for immunohistochemistry are listed in Supplementary Table 8.

### In situ hybridization analysis

Digoxigenin (DIG) or fluorescein labeled RNA probes were made using the DIG RNA labeling kit (Roche, Cat. 11175025910) or fluorescein RNA labeling kit (Roche, cat. no. 11685619910), respectively, per manufacturer’s protocol. T7 RNA polymerase (Roche, cat. no. 10881767001), T3 RNA polymerase (Roche, cat. no. 11031171001) or SP6 RNA polymerase (Roche, cat. no. 10810274001) were employed to generate RNA, depending on the particular RNA probe. After DNase I treatment and precipitation, RNA probes were dissolved in 40 μl RNAse-free water. Forelimbs or hindlimbs were dissected and fixed in 4% paraformaldehyde (made with DEPC-treated PBS) at 4°C overnight, washed with PBS, and incubated in 30% sucrose (made with DEPC-treated H2O) at 4°C overnight. Tissues were embedded in OCT and frozen sections were cut at 20 μm using a cryostat. Sections were fixed again with 4% paraformaldehyde (made with DEPC-treated PBS) at room temperature for 5 minutes, washed twice (5 minutes each at room temperature) with PBS containing 0.1% Tween 20 (PBST). The sections were digested with Proteinase K (4 μg/ml in PBS) at room temperature for 8-20 minutes (depending on the RNA probe), washed twice with PBST (5 minutes each) and fixed again with 4% paraformaldehyde (in PBS) at room temperature for 5 minutes. After washing twice with PBST (5 minutes each, at room temperature), sections were acetylated for 10 minutes (at room temperature) in a solution containing 0.25% acetic anhydride and 0.1M triethanolamine (in DEPC-treated H2O). After washing twice with PBST (5 minutes each, at room temperature) the slides were rinsed in DEPC-treated H2O, and then air dried. The RNA probe (listed in Supplemental Table 10) was diluted to 100 ng/ml in a 200 μl volume of hybridization solution (as described in ^72^), and heated at 95 °C for 5 minutes before adding to the slide. Hybridization was performed at 65 °C overnight, and washed according to the protocol outlined in ^72^. After hybridization, washes, and blocking endogenous peroxidase activity, the slides were blocked with 10% heat inactivated goat serum at room temperature for one hour. Anti-DIG-POD antibody (1:100 dilute, Anti-Digoxigenin-POD, Fab fragment, 11207733910, Roche) or Anti-Fluorescein-POD antibody (1:100, Anti-Fluorescein-POD, Fab fragments, 11426346910, Roche) was added and the slides were incubated at 4 °C overnight. After washing in TNT Buffer (100 mM Tris–HCl pH 7.5, 150 mM NaCl, 0.05% Tween20), the biotin (1:100) amplification reagent (TSA Biotin Kit, NEL749A001KT) or Fluorescein (1:100) amplification reagent (TSA plus Cyanine 3/ Fluorescein System, NEL753001KT) was applied and slides incubated 20 minutes at room temperature to develop the signal. Biotin signal was further detected by Streptavidin Secondary Alexa 594 dyes. Slides were washed in TNT Buffer (3 × 10 minutes at room temperature), rinsed in H2O, and mounted under a coverslip with Aqua-mount (13800; Lerner Labs).

### Statistics and Reproducibility

RNA-Seq was performed with duplicate biological samples. Differential expression of genes between sample types in these data sets was determined through the Exact Test in edgeR, with Benjamini-Hochberg ^68^ multiple testing correction (FDR) set at < 0.05. ATAC-Seq was performed with two biological repeats to ensure the robustness of the data sets. RT-qPCR assays and luciferase assays were performed with 2-3 biological repeats and each assay was performed with technical repeats. Prism statistical software (GraphPad) was employed to analyze the data. Statistical analysis was performed using either unpaired or paired t tests.

### RNA-Seq and ATAC-Seq primary data

All primary RNA-Seq and ATAC-Seq data sets have been deposited with GEO.

To review GEO accession GSE132379: Go to https://www.ncbi.nlm.nih.gov/geo/query/acc.cgi?acc=GSE132379 Enter token sfgtgoakdvidpaf into the box

## Acknowledgments

We thank Attila Aszodi, Veronique Lefebvre, Maurizio Pacifici and Cliff Tabin for providing in situ probes; Terence Capellini, April Craft, Vicki Rosen, Matt Warman and Yingzi Yang for their constructive comments on the manuscript; and the Nikon Imaging Center at Harvard Medical School for use of their microscopes and cameras. We thank Mary Goldring for supplying us with human costal chondrocyte cell lines, and Nicholas O’Neill and Shariq Madha for assistance with computational analyses. We acknowledge the use of tissues procured by the National Disease Research Interchange (NDRI) with support from NIH grant U42OD11158. Portions of this research were conducted on the O2 High Performance Compute Cluster, supported by the Research Computing Group, at Harvard Medical School.

## Author Contributions

C.-H. Z. and A. B. L. designed experiments and wrote the paper. C.-H. Z. and Y. G. conducted experiments and edited the paper. U. J. provided experimental expertise, analyzed RNA-Seq and ATAC-Seq data, and edited the paper. H.-H. H. isolated superficial and deep zone bovine articular cartilage tissue and edited the paper. K. M. H. performed bioinformatics analysis of the RNA-Seq data. A. J. G. and R. A. S. provided experimental expertise and edited the paper.

## Data and correspondence

All primary RNA-Seq and ATAC-Seq data sets have been deposited with GEO (GEO accession GSE132379). All data, code, and materials used in the analysis will be made available to any researcher for purposes of reproducing or extending the analysis. Correspondence and material requests should be addressed to A. B. L.

## List of Supplementary Materials

### Supplementary Figure Legends

**Supplementary Figure 1.**
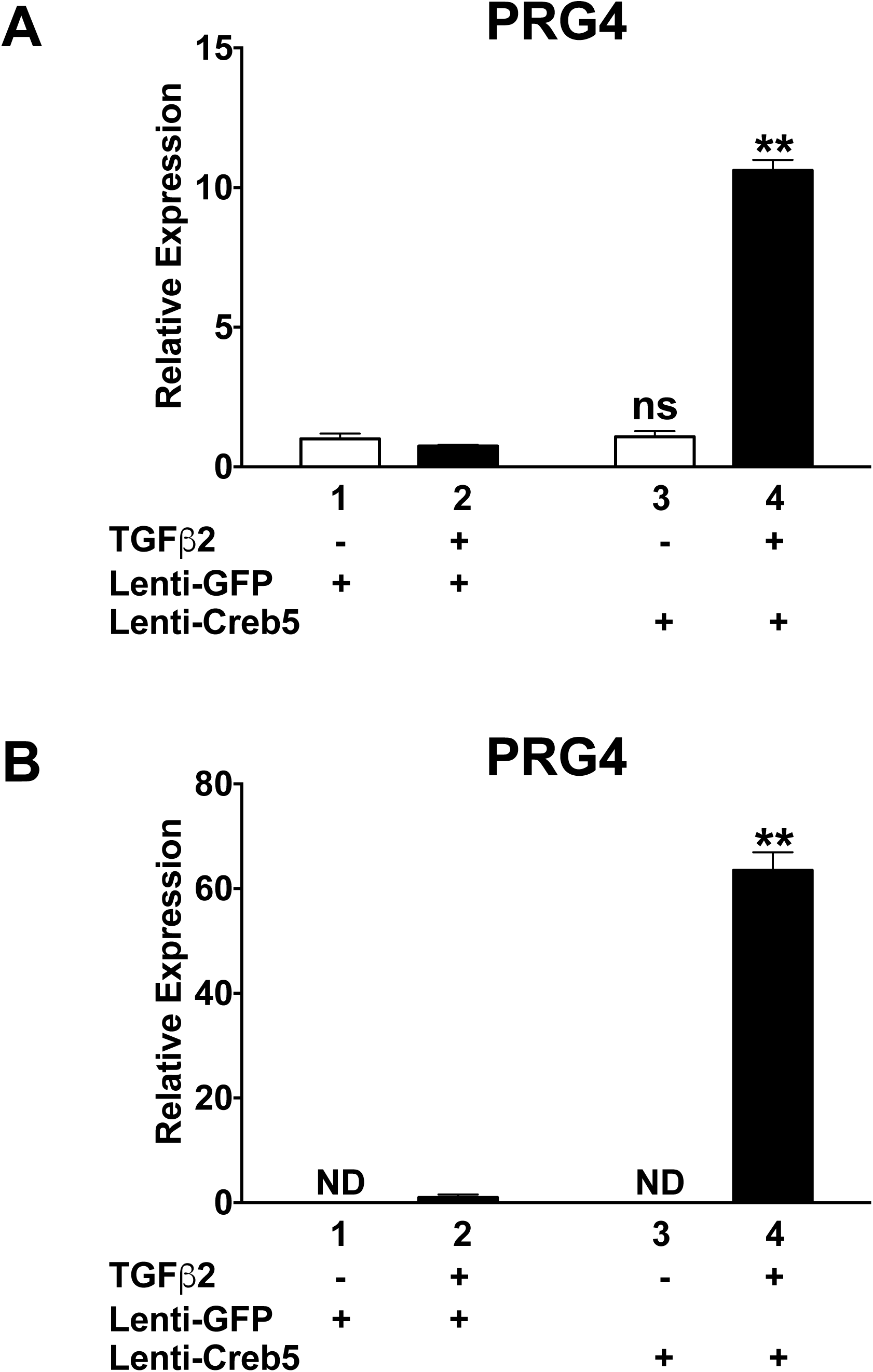
Exogenous Creb5 can promote TGF-β dependent induction of Prg4 expression in either a human chondrosarcoma cell line (SW1353) or in an immortalized human costal chondrocyte cell line (C-28/I2; ^49^). (A) Sw1353 human chondrosarcoma cells (ATCC HTB 94) or (B) C-28/I2 cells were infected with control lentivirus (Lenti-GFP) or Lenti-Creb5. After selection in puromycin, the cells were cultured in either the presence or absence of TGF-β2 (20 ng/ml) for 3 days in ultra-low attachment plates. Gene expression was assayed by RT-qPCR and normalized to *GAPDH*. Lanes 3 and 4 are compared to lanes 1 and 2, respectively. *P<0.05, **P<0.01, ***P<0.001, ND (not detected) and ns (non-significant difference) are indicated and error bar indicates standard error of the mean. Similar results have been obtained in 3 independent biological repeats.

**Supplementary Figure 2.**
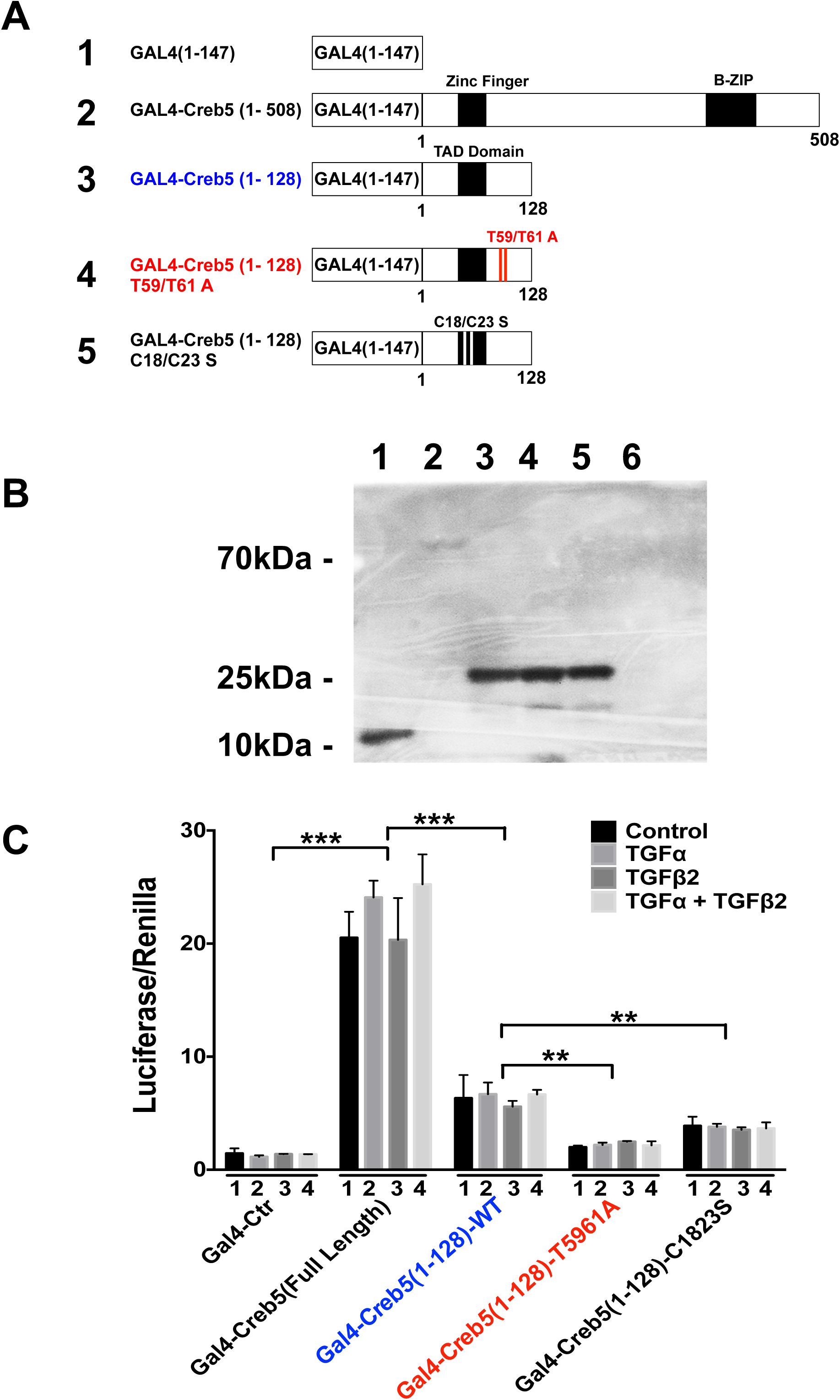
The amino-terminus of Creb5 contains a relatively weak transcriptional activation domain, which requires both SAPK phosphorylation sites and an intact zinc finger domain for maximal activity. (A) Diagrams of GAL4 DNA binding domain fusions with either full length Creb5 (1-508) or with the N-terminus of Creb5 (1-128). Mutations in either the SAPK phosphorylation sites (T59/T61A) or in the zinc finger domain (C18/C23S) are displayed. (B) Expression of various GAL4 DNA binding domain fusions (as numbered and diagrammed in A) in transfected 293T cells, as detected by a GAL4 western blot. Lane 6 is derived from non-transfected 293T cells. (C) Superficial zone bovine articular chondrocytes were transfected with various GAL4-Creb5 constructs, plus a GAL4-firefly luciferase reporter and a CMV-renilla luciferase reporter; cells were cultured in either the absence or presence of TGF-β2 (20 ng/ml) and TGF-α (100 ng/ml), as indicated for 3 days. Relative expression of firefly/renilla luciferase is displayed. The relative level of GAL4-firefly luciferase/renilla luciferase, in response to co-transfection of the various GAL4 constructs, were compared across the 4 conditions by paired t test, as indicated. **P<0.01, ***P<0.001 are indicated and error bar indicates SEM.

**Supplementary Figure 3.**
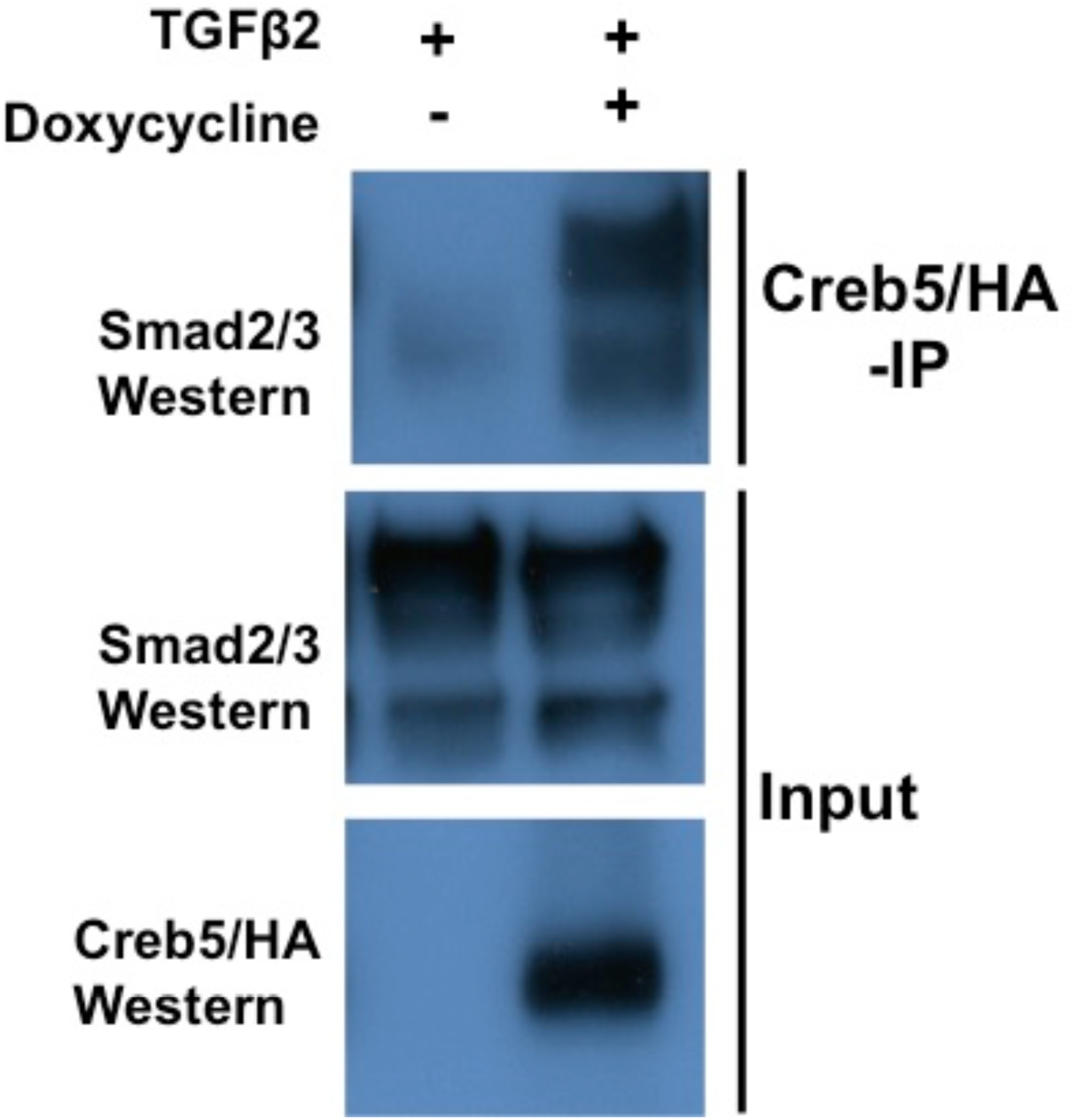
Creb5 can co-immunoprecipitate Smad2/3. Bovine deep zone articular chondrocytes that were programmed to express doxycycline-inducible Creb5 lentivirus (DZC-iCreb5-HA) were cultured in ultra-low attachment dishes in the presence of TGF-β2 (20 ng/ml) with or without doxycycline (1μg/ml) for 3 days. After 3 days culture, cells were collected and co-immunoprecipitation was performed using monoclonal anti-HA agarose. Immunoprecipitated protein was detected using anti-Smad2/3 antibody in western blot. Input protein was detected using anti-Smad2/3 and anti-HA antibody.

### Supplementary Tables

**Supplementary Table 1.**
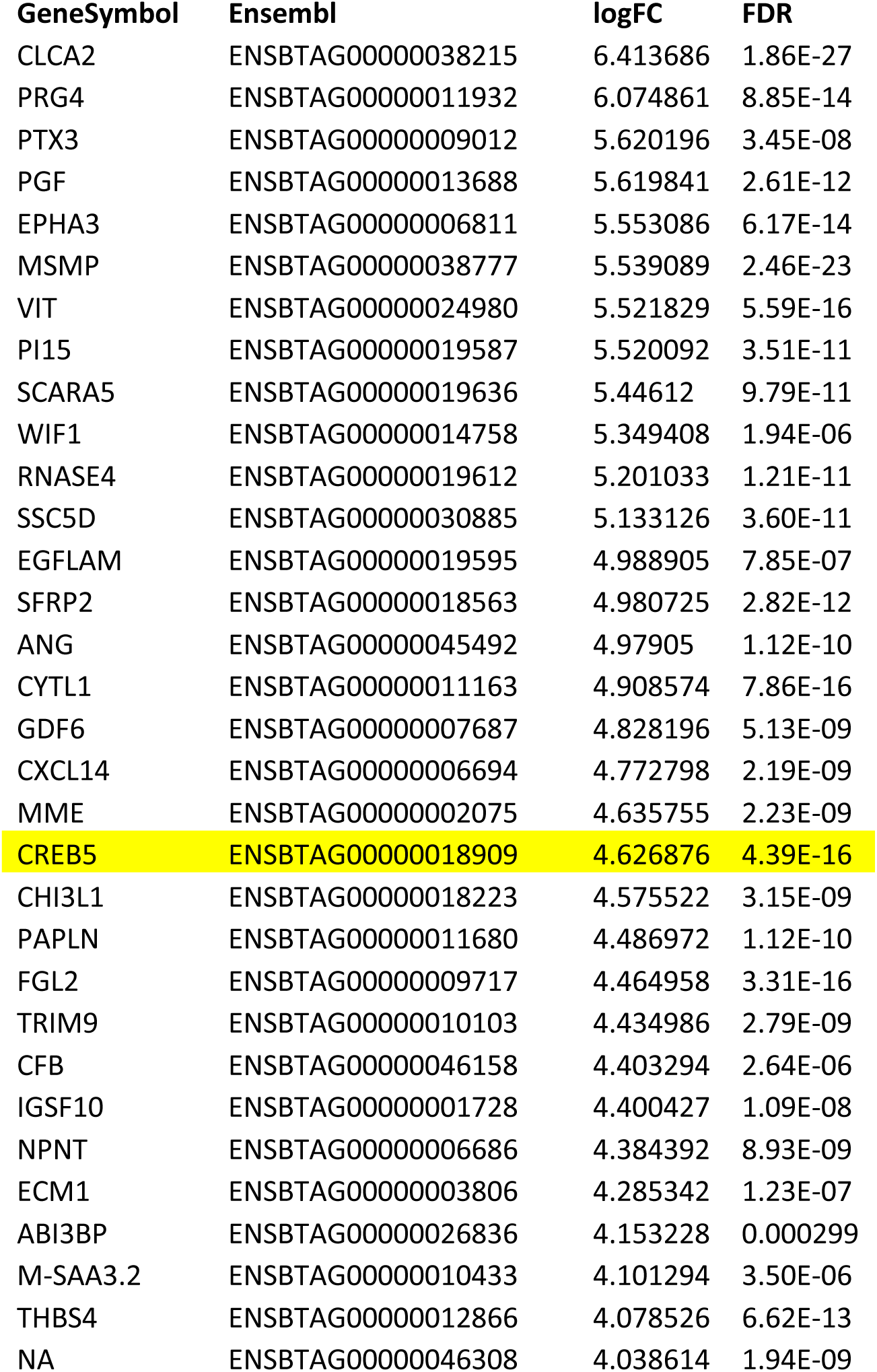
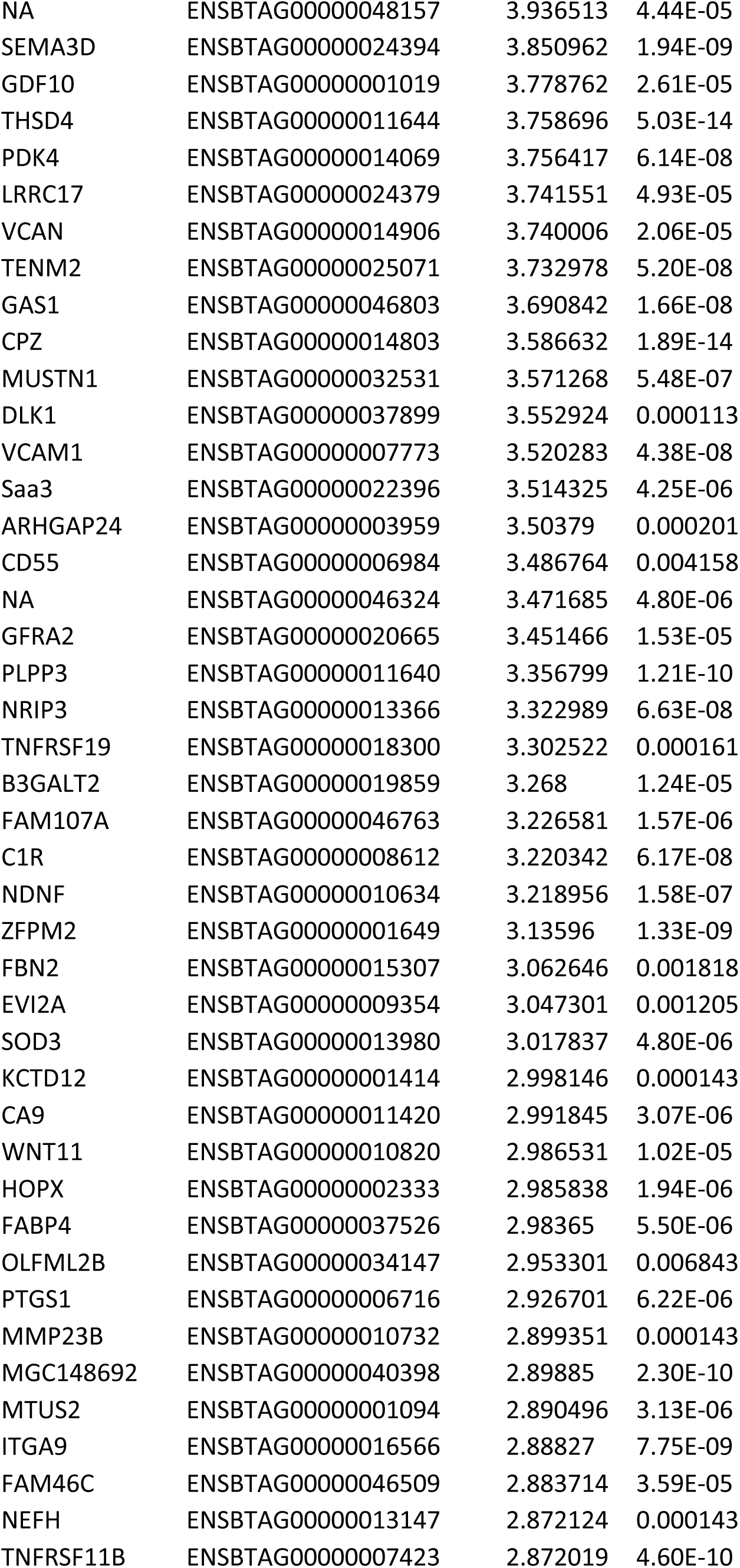
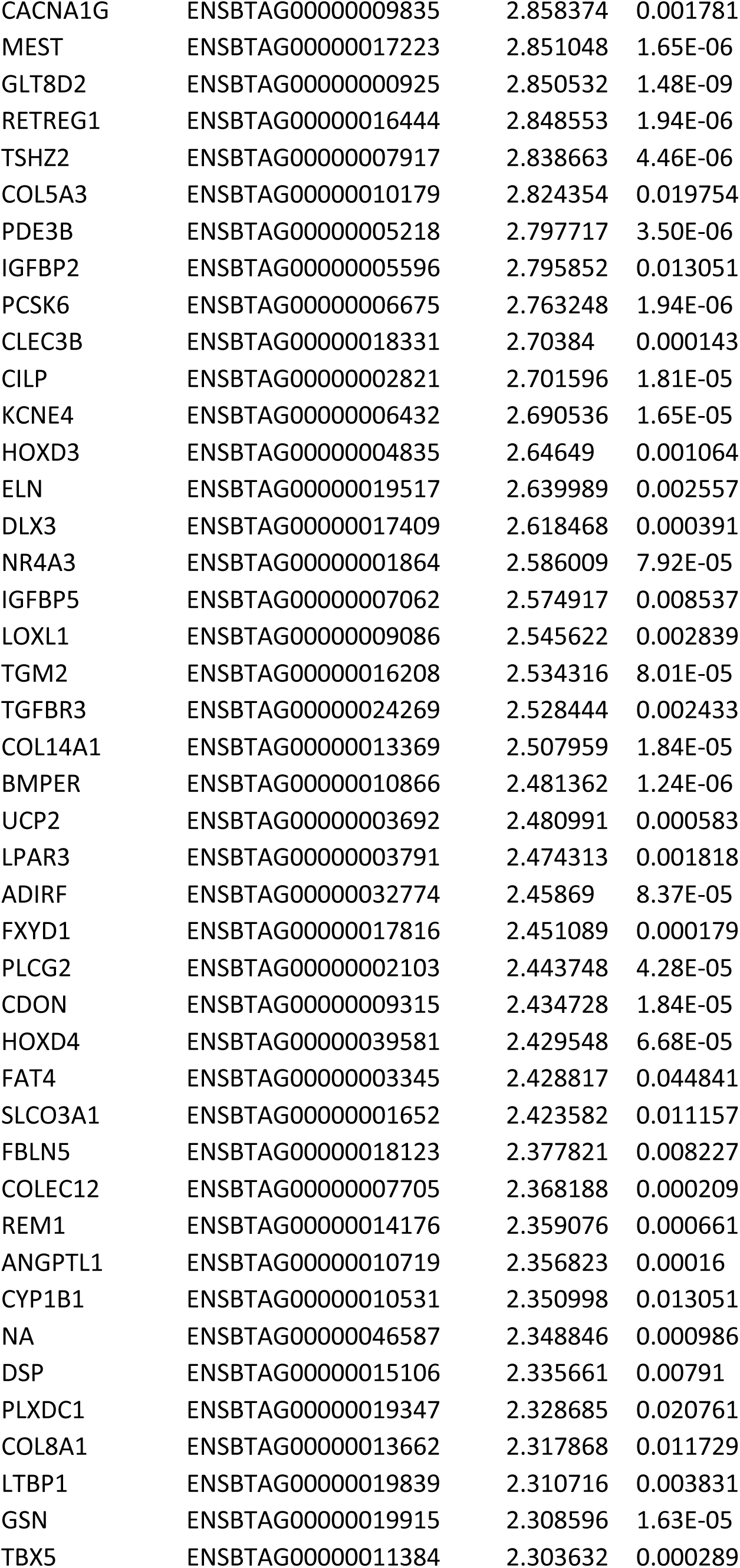
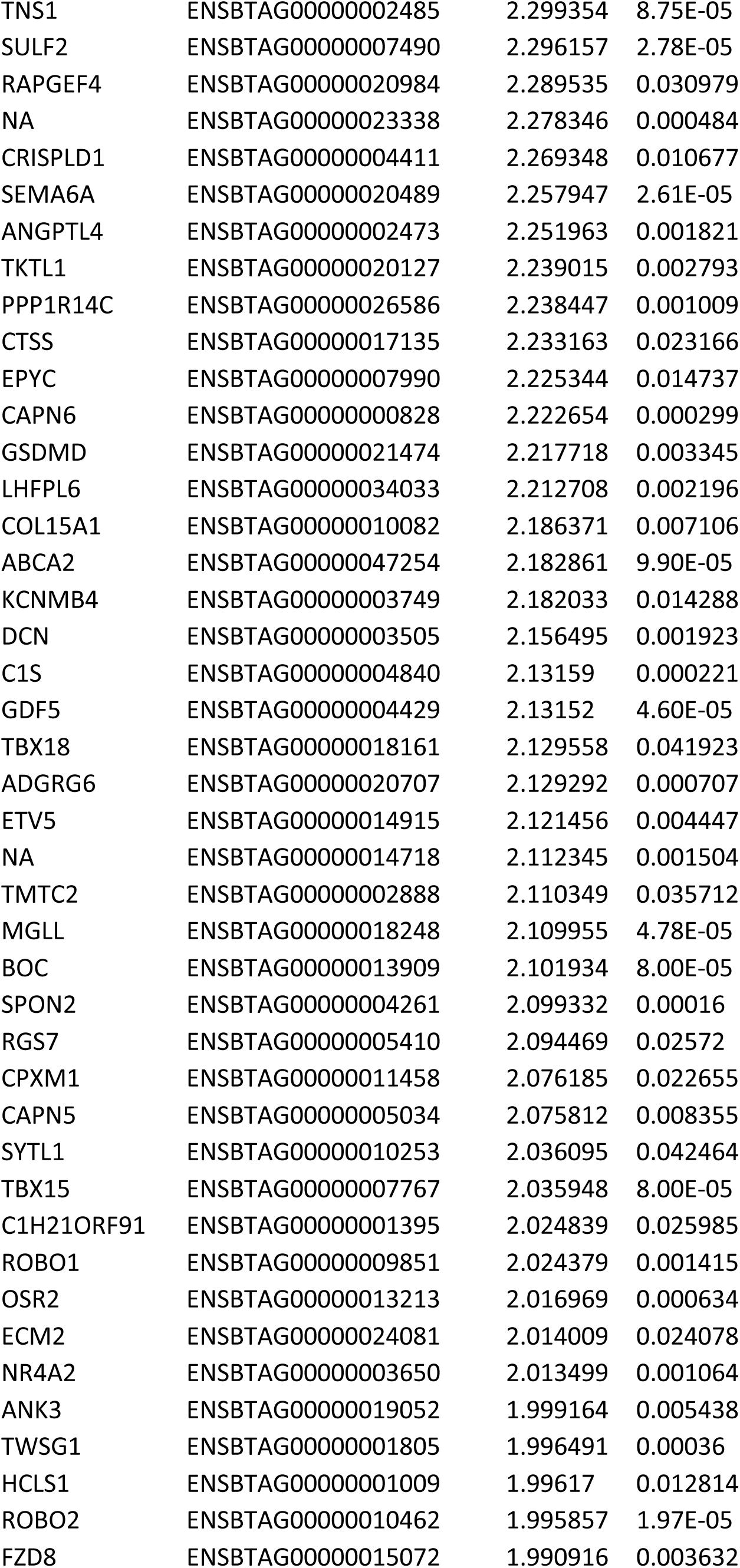
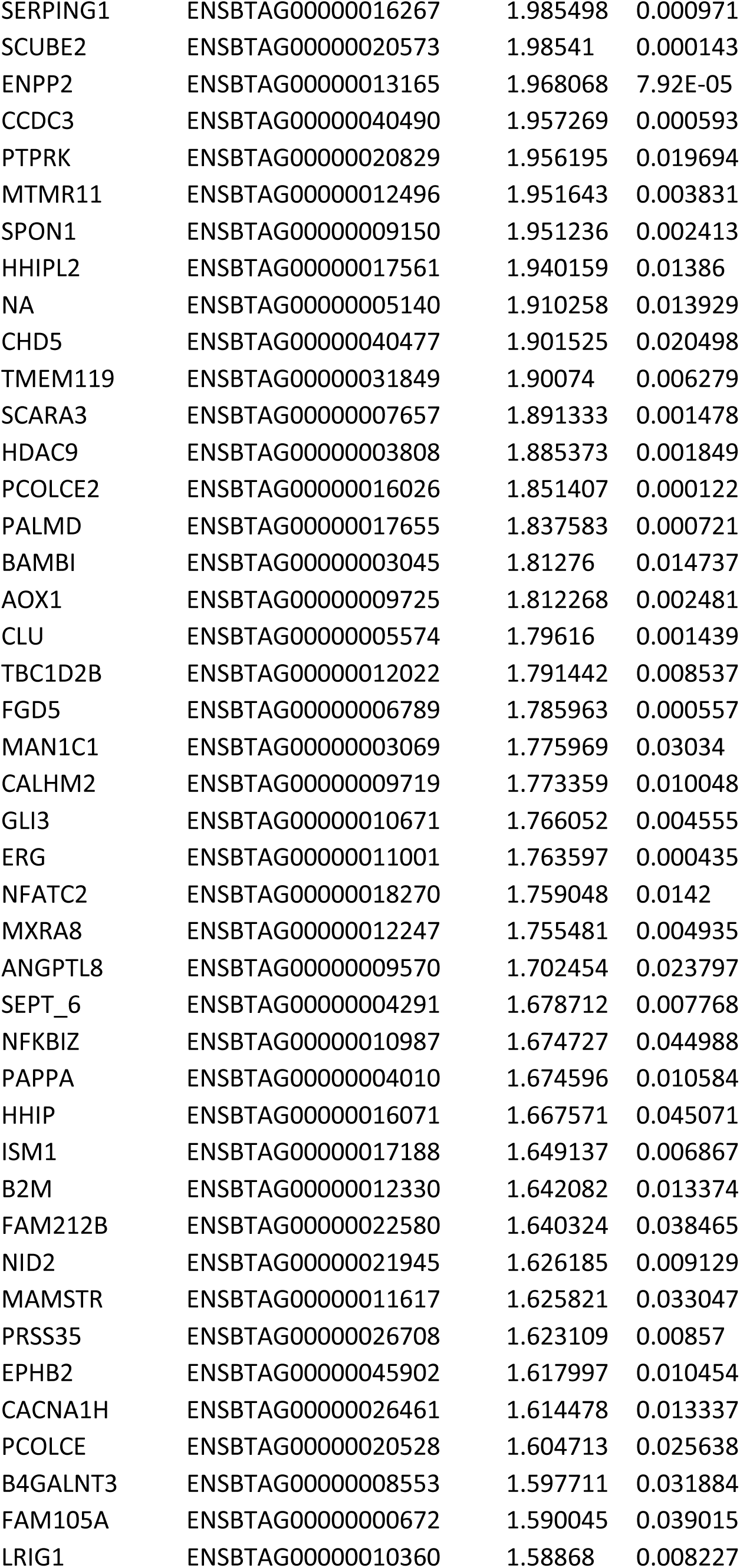
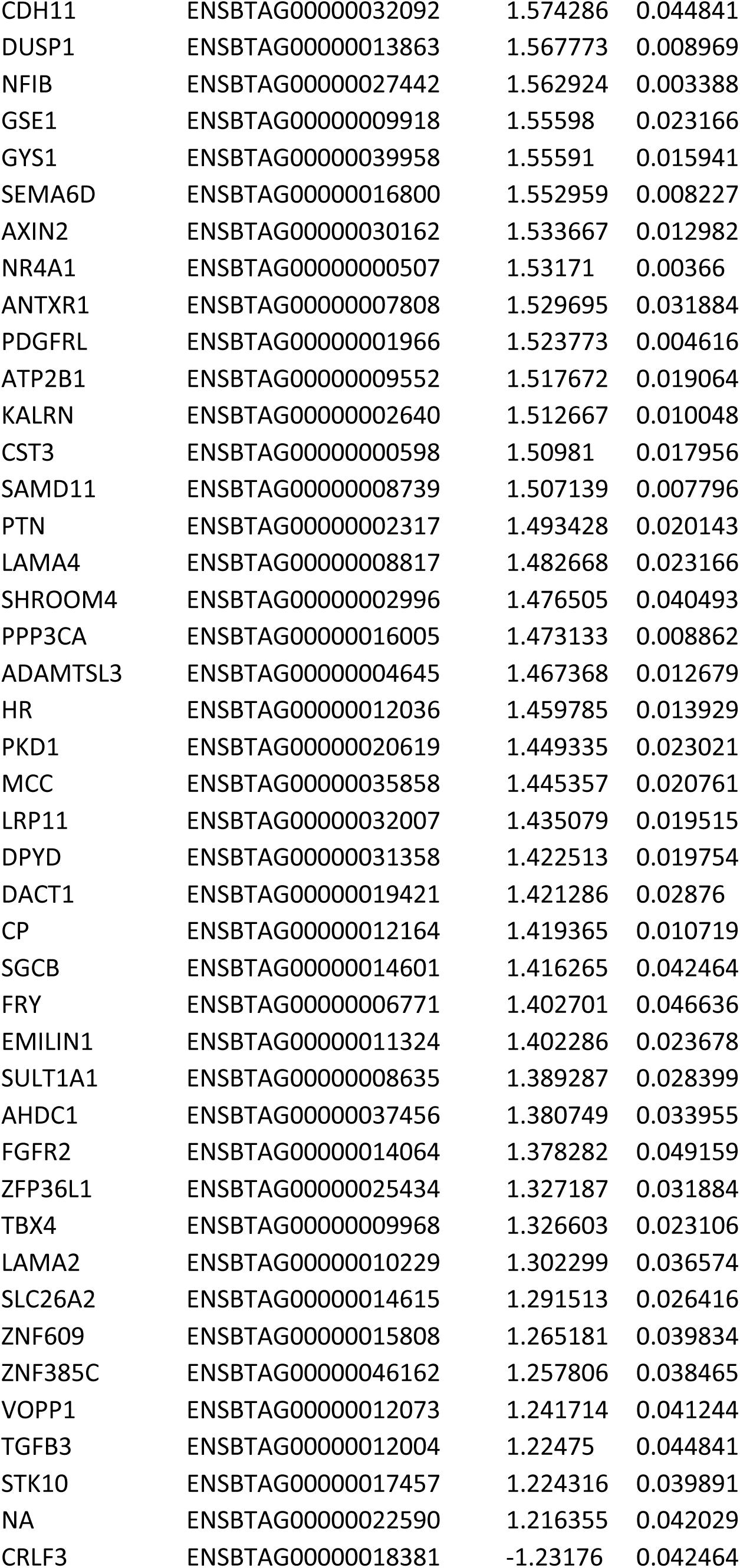
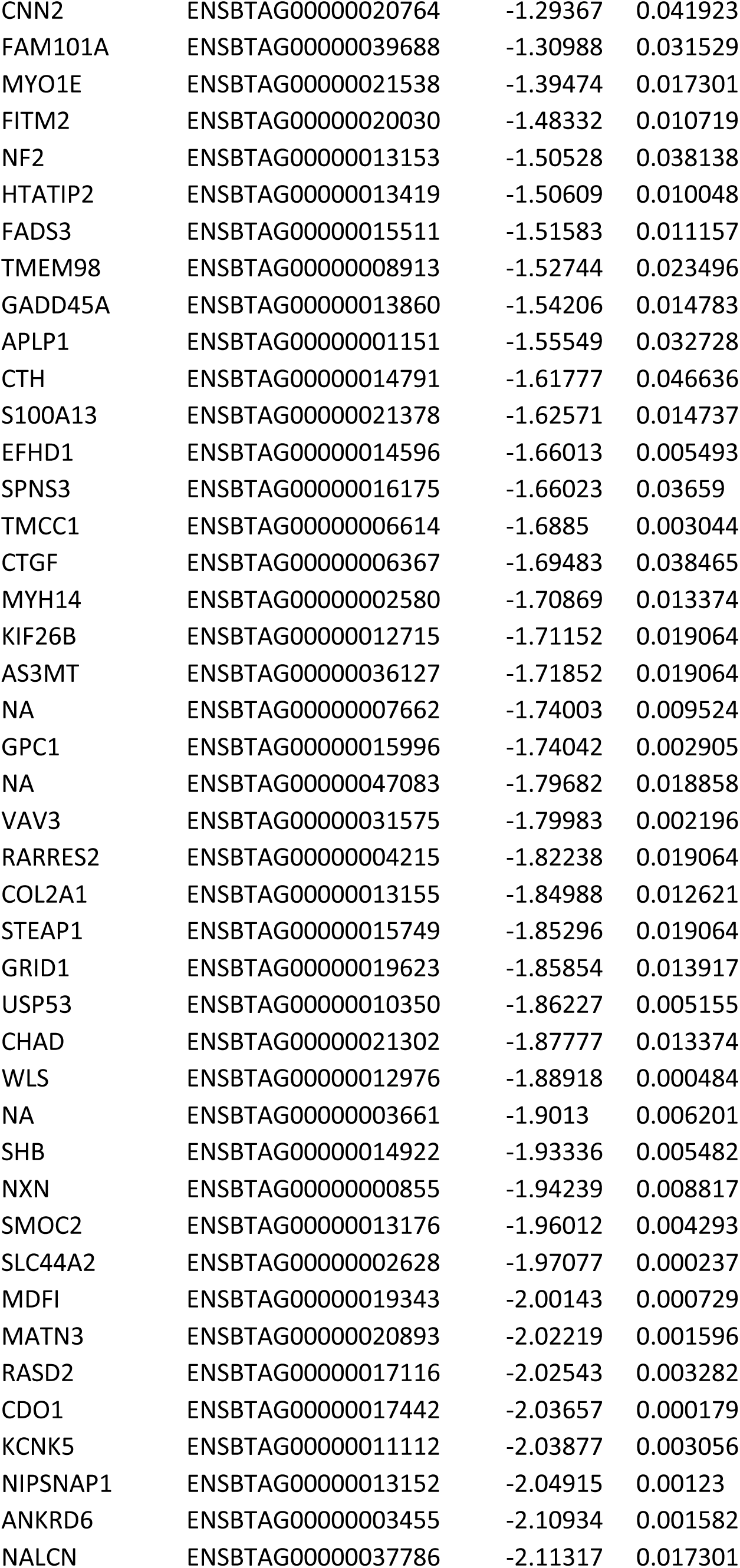
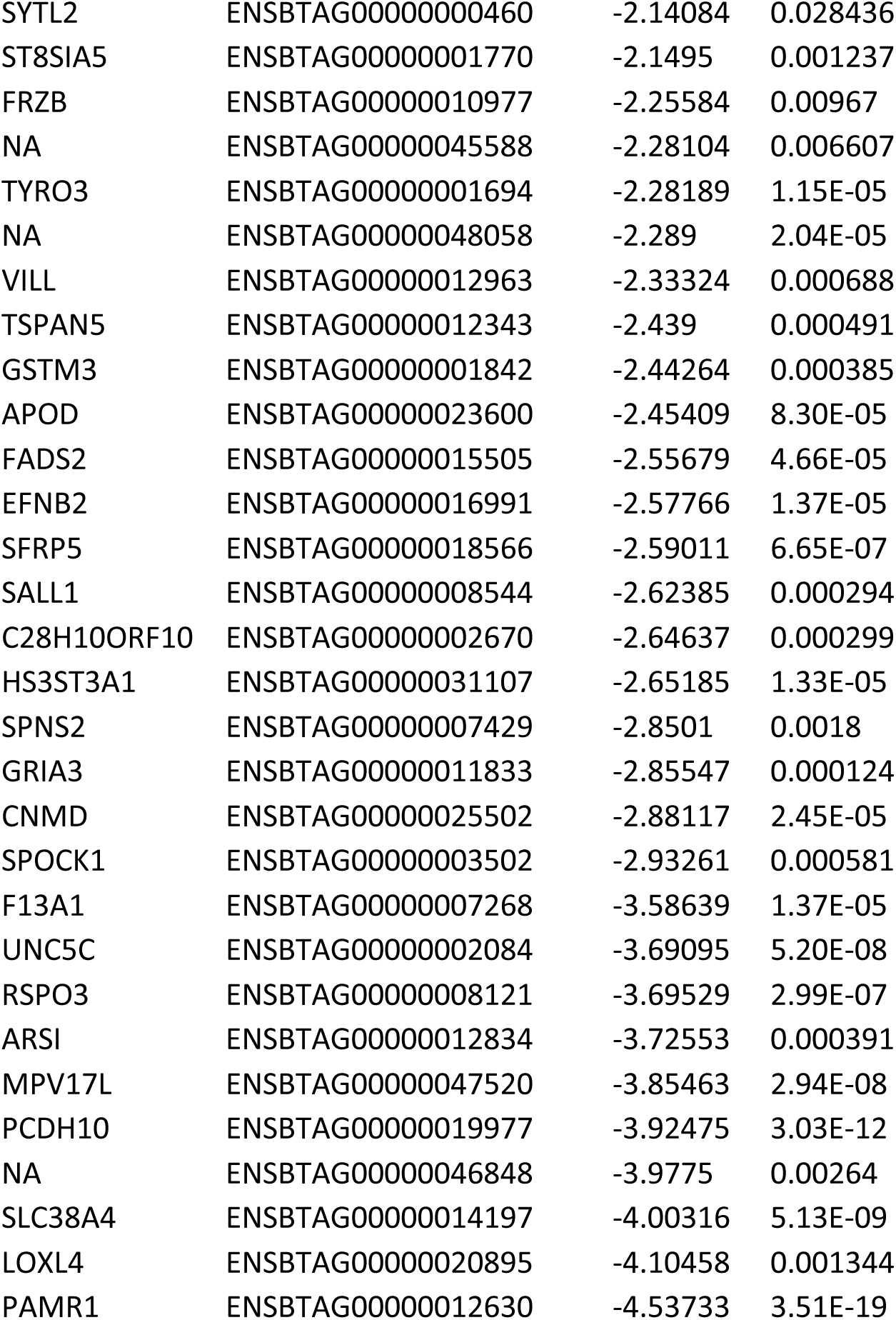
List of genes that are differentially expressed in superficial versus deep zone bovine articular chondrocytes. 320 genes are listed whose expression differed (by indicated LogFC) between SZC and DZC (false discovery rate, FDR <0.05).

**Supplementary Table 2:**
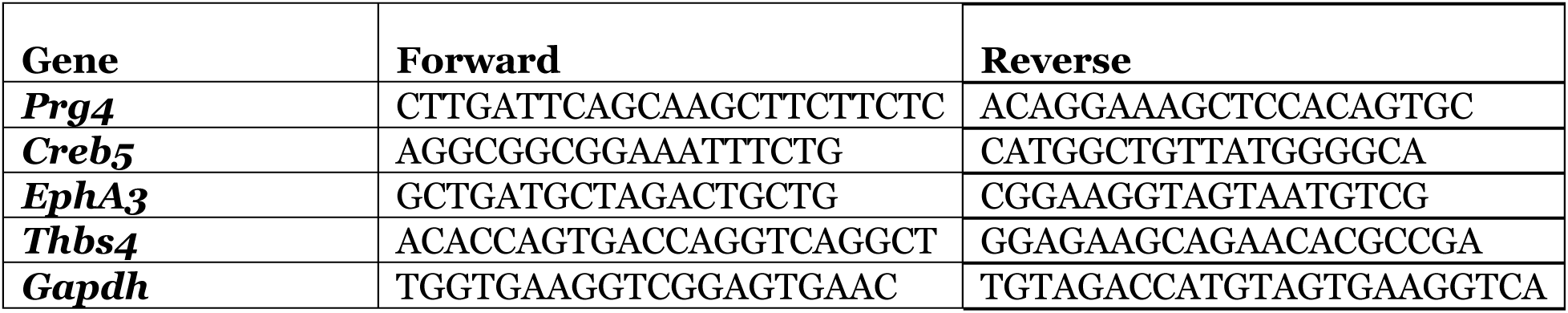
RT-qPCR Primers to amplify bovine cDNA.

**Supplementary Table 3:**
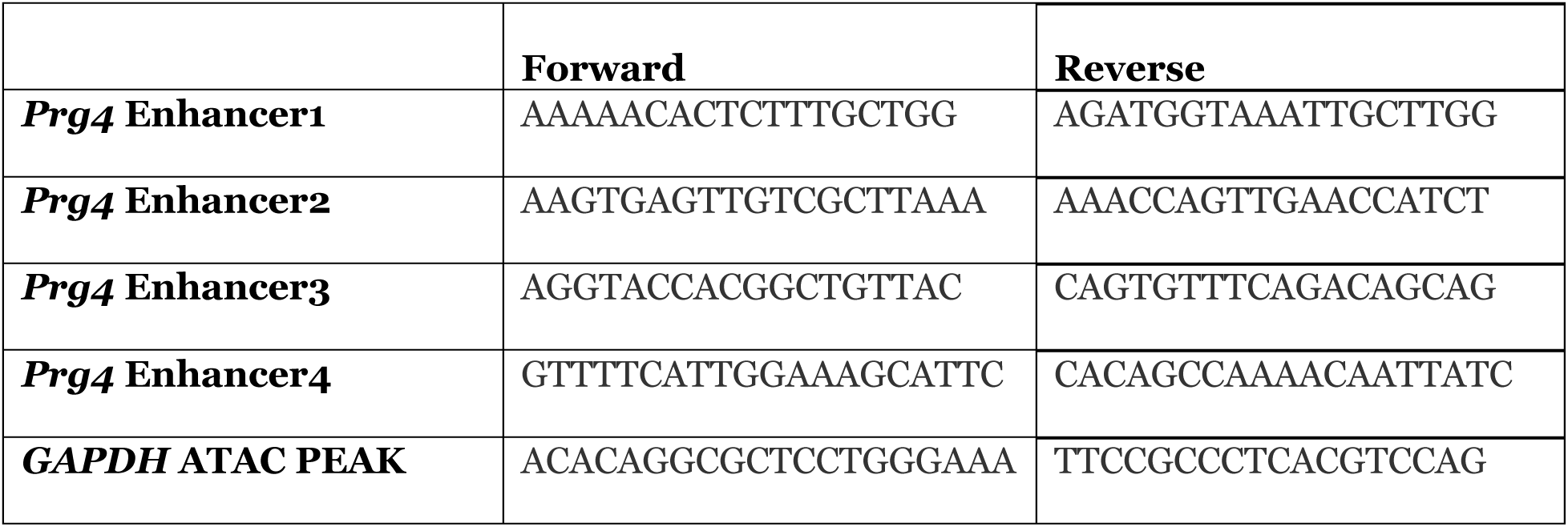
ChIP-PCR Primers.

**Supplementary Table 4:**
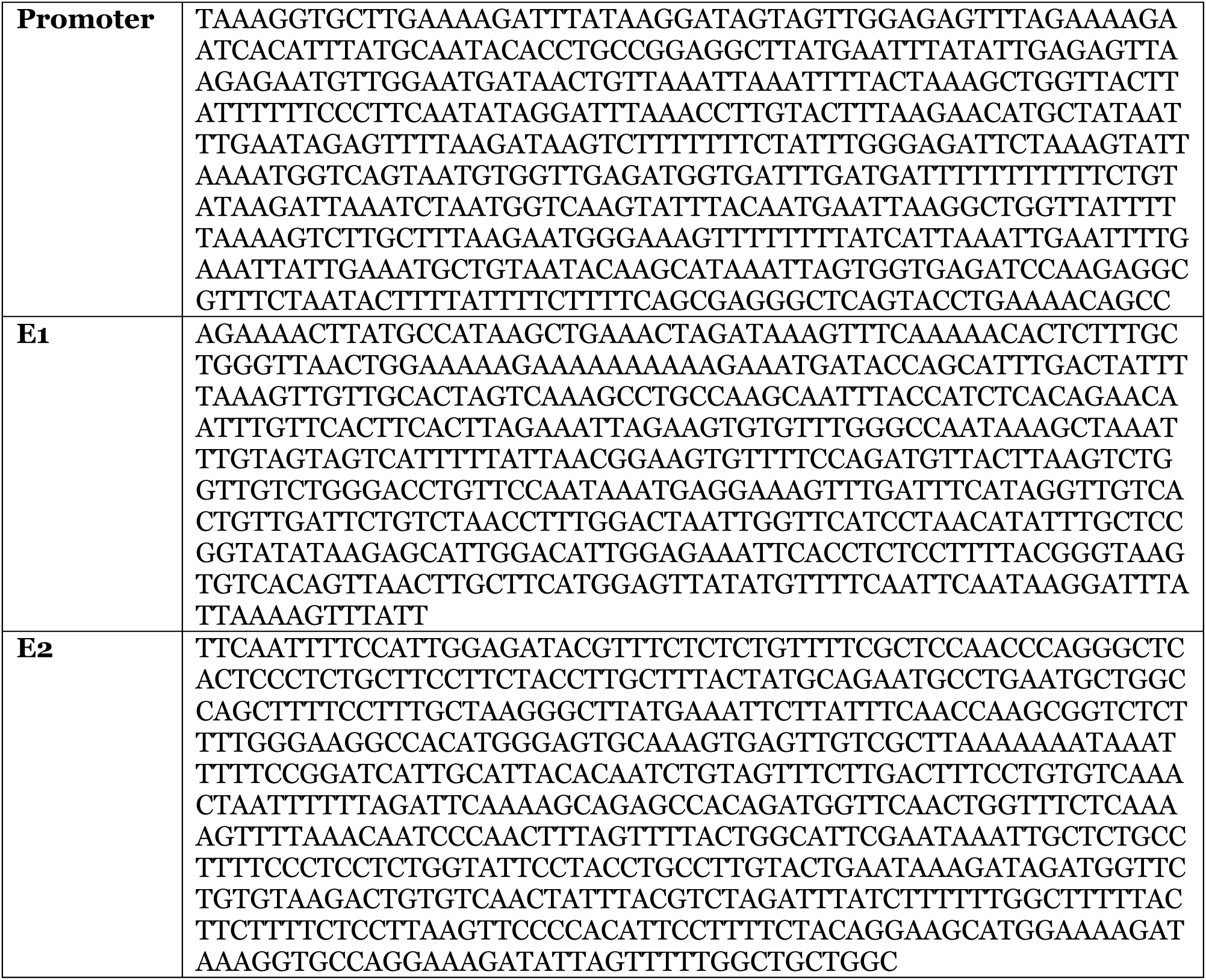
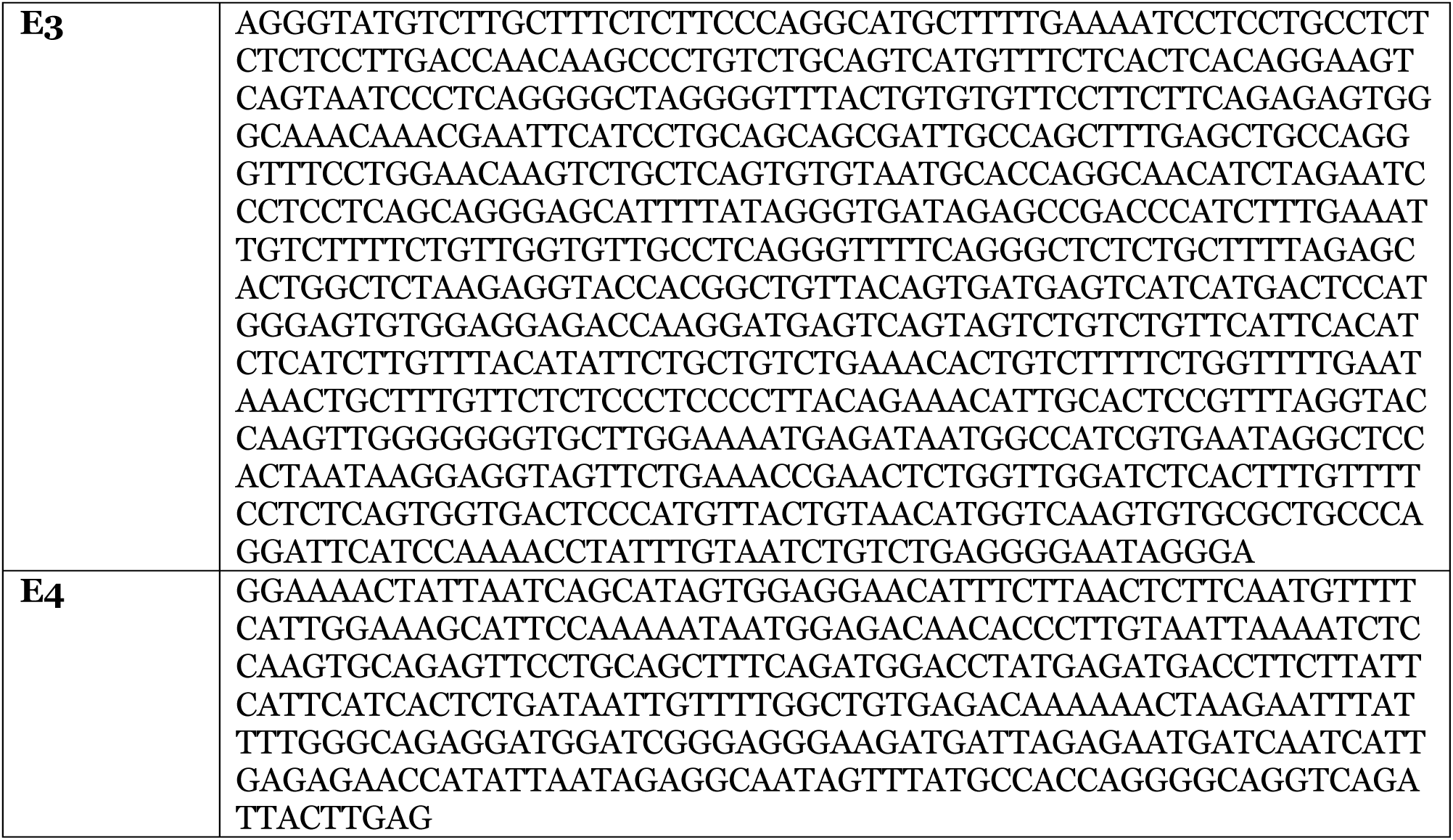
Prg4 Promoter and Enhancer E1,E2,E3,E4 sequences.

**Supplementary Table 5:**
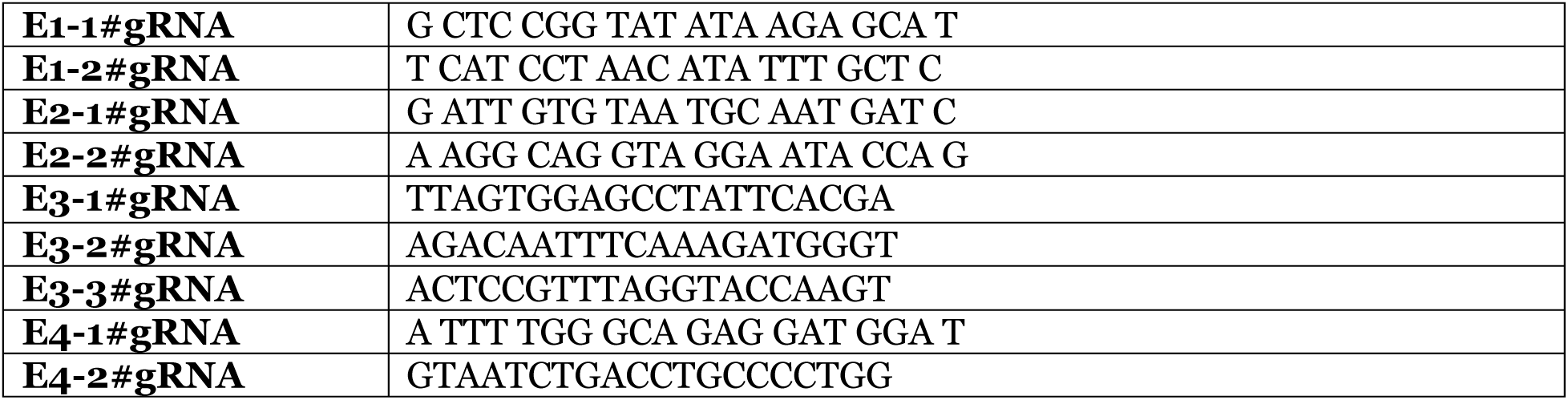
Sequence of dCas9-KRAB guide RNAs that target the bovine Prg4 enhancers.

**Supplementary Table 6:** Antibodies employed for Western Blots.

| Cat. No. | Concentration | Target | Host | Vendor |
| --- | --- | --- | --- | --- |
| PA5-65593 | 1:1000 | Creb5 | Rabbit | Thermo Fisher |
| 9221S | 1:500 | p-Creb5 | Rabbit | Cell Signaling Technology |
| 9211S | 1:1000 | p-p38 | Rabbit | Cell Signaling Technology |
| 9212S | 1:1000 | P38 | Rabbit | Cell Signaling Technology |
| T9026 | 1:1000 | α-Tubulin | Mouse | Sigma |
| ab9110 | 1:1000 | HA | Rabbit | abcam |
| sc-805 | 1:1000 | HA | Rabbit | Santa cruz biotechnology |
| A14635 | 1:1000 | Creb5 | Rabbit | ABclonal |
| sc-510 | 1:500 | GAL4 | Mouse | Santa cruz biotechnology |
| 8685S | 1:1000 | Smad2/3 | Rabbit | Cell Signaling Technology |

**Supplementary Table 7:** Antibodies employed for either ChIP or co-immunoprecipitation (Co-IP)

| Cat. No. | Concentration | Target | Host | Vendor |
| --- | --- | --- | --- | --- |
| ab9110 | 5 μg per ChIP | HA | Rabbit | abcam |
| ab46540 | 5 μg per ChIP | IgG | Rabbit | abcam |
| A2095 | 25 μl per Co-IP | HA | Mouse | Sigma |

**Supplementary Table 8:**
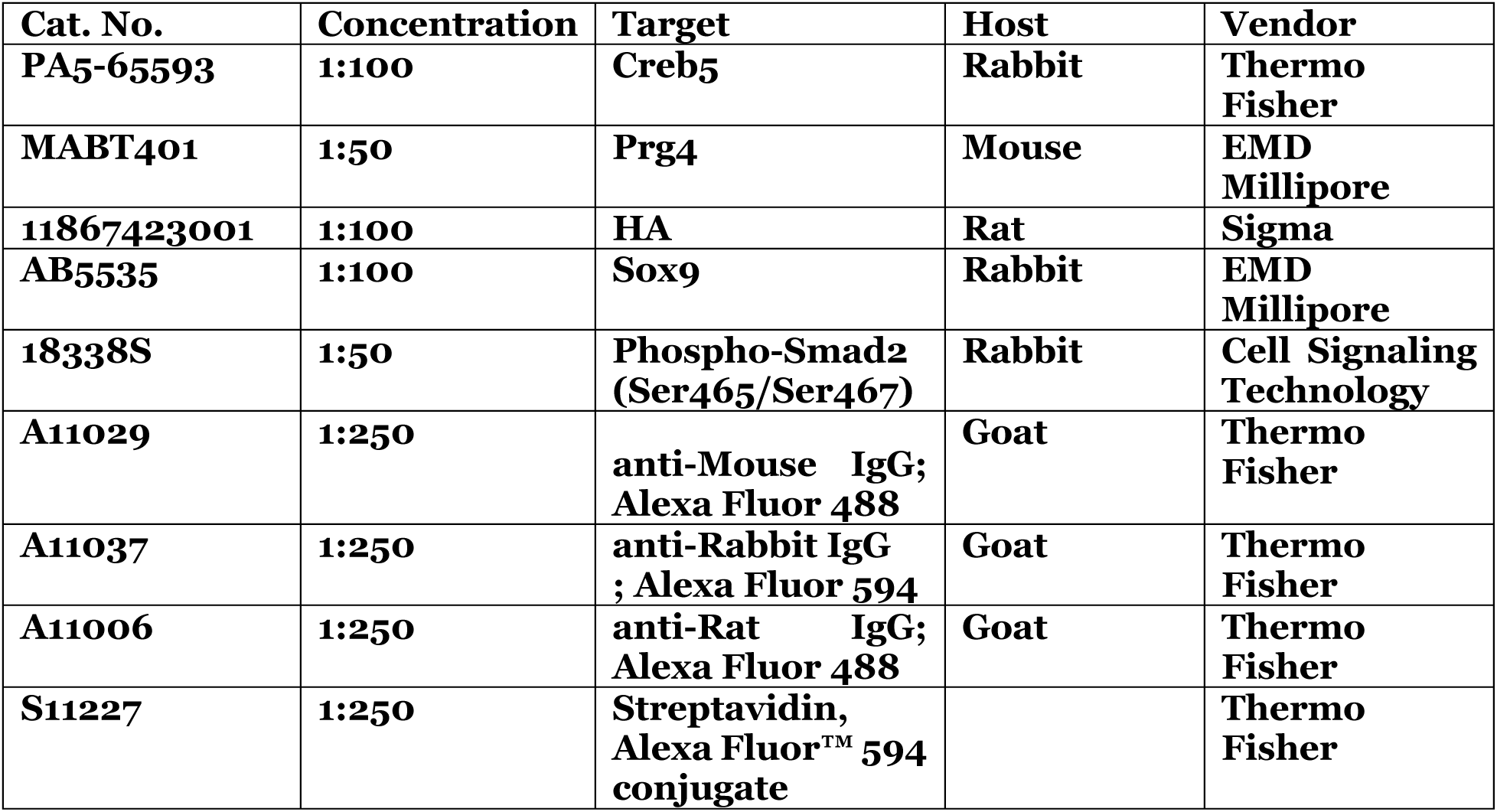
Antibodies employed for immunohistochemistry.

**Supplementary Table 9:** Growth factors and inhibitors used for chondrocyte culture.

|  | **Cat. No.** | **Concentration** | **Vendor** |
| --- | --- | --- | --- |
| **TGF-α** | **239-A-100** | **100 ng/ml** | **R&D Systems** |
| **TGF-β2** | **302-B2-010** | **20 ng/ml** | **R&D Systems** |
| **Doxycycline** | **D9891-1G** | **1 μg/ml** | **Sigma** |
| **SB203580** | **152121-47-6** | **10 μM** | **TOCRIS** |
| **SP600125** | **129-56-6** | **10 μM** | **TOCRIS** |

**Supplementary Table 10:**
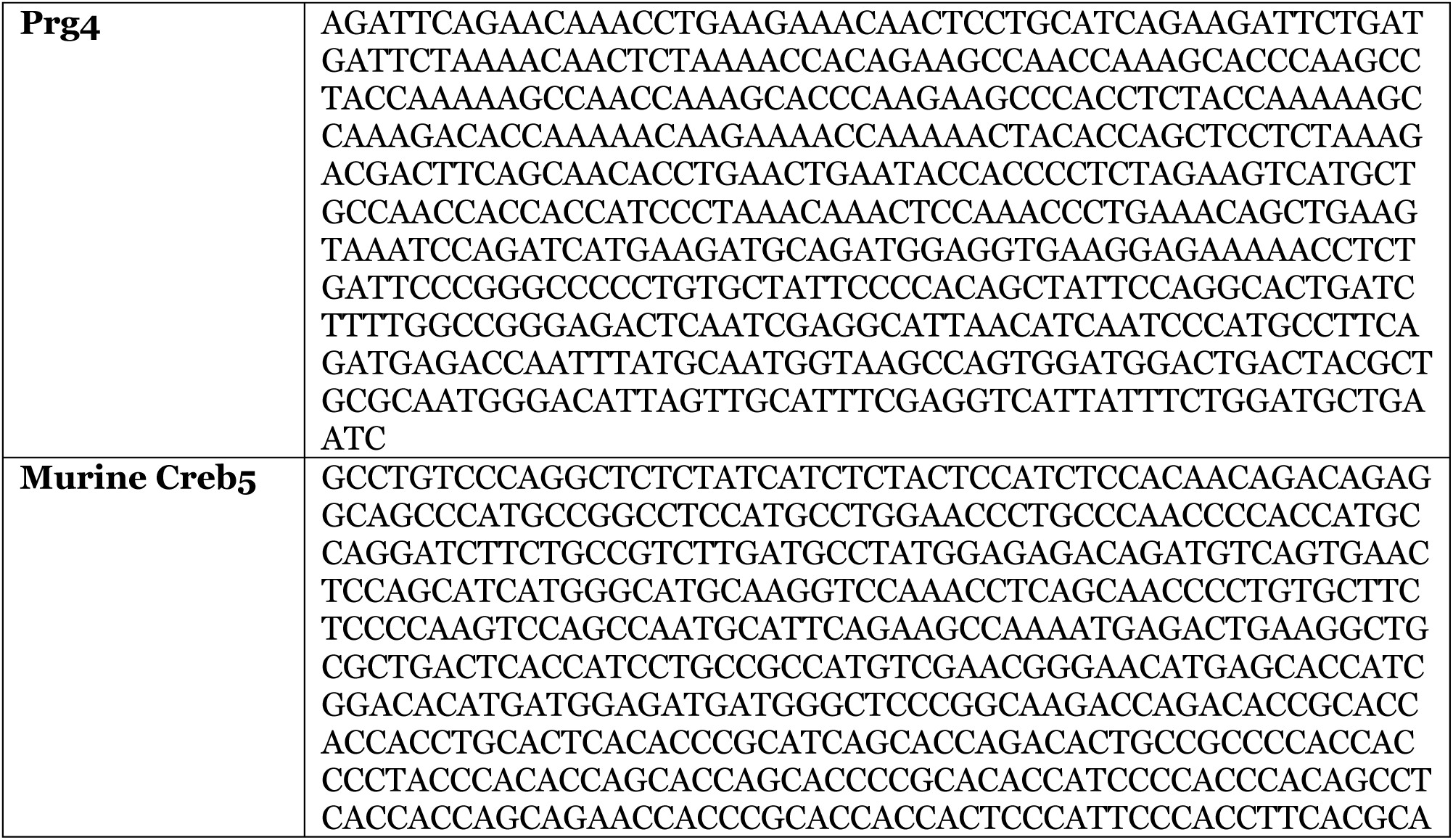
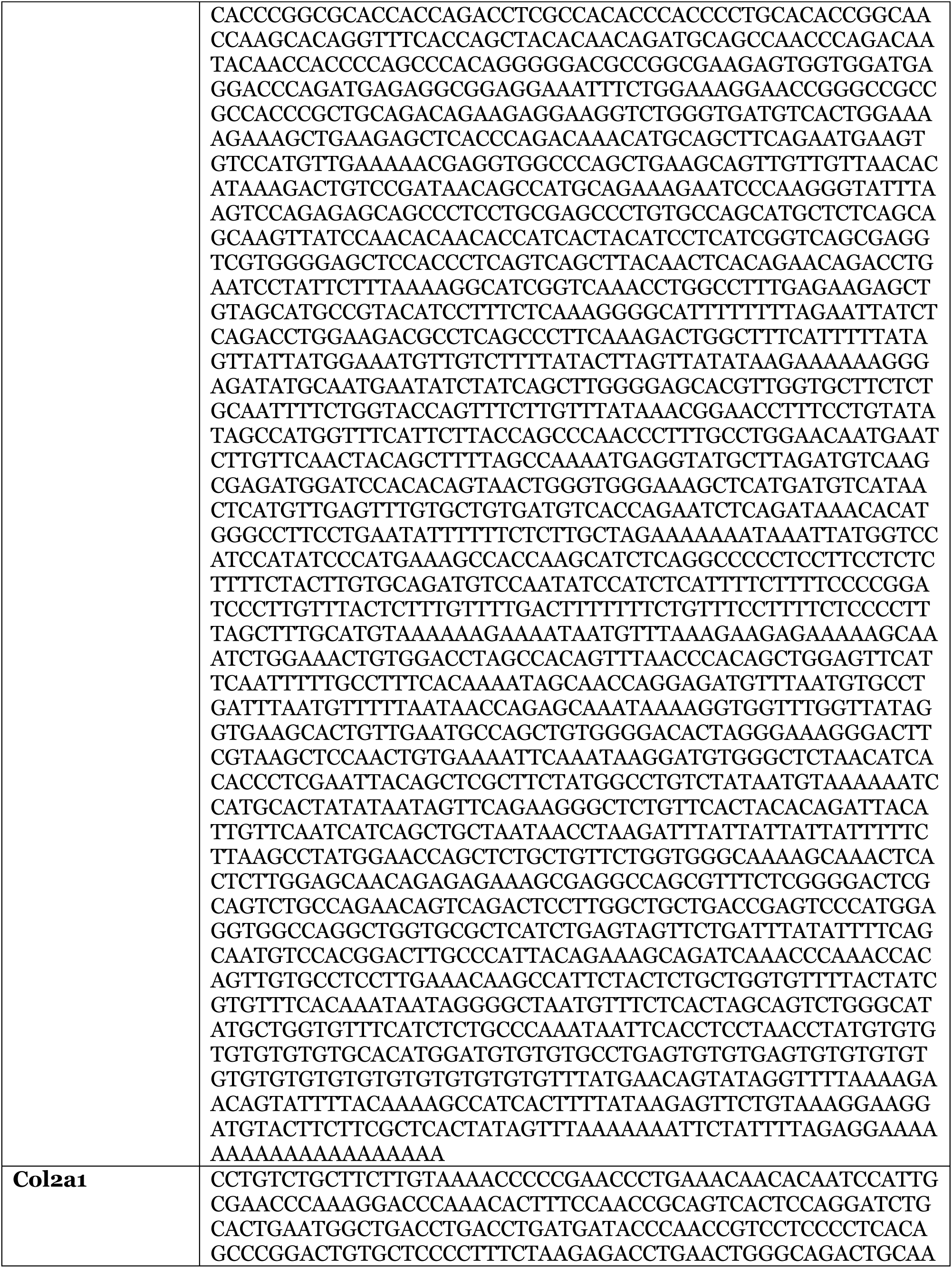
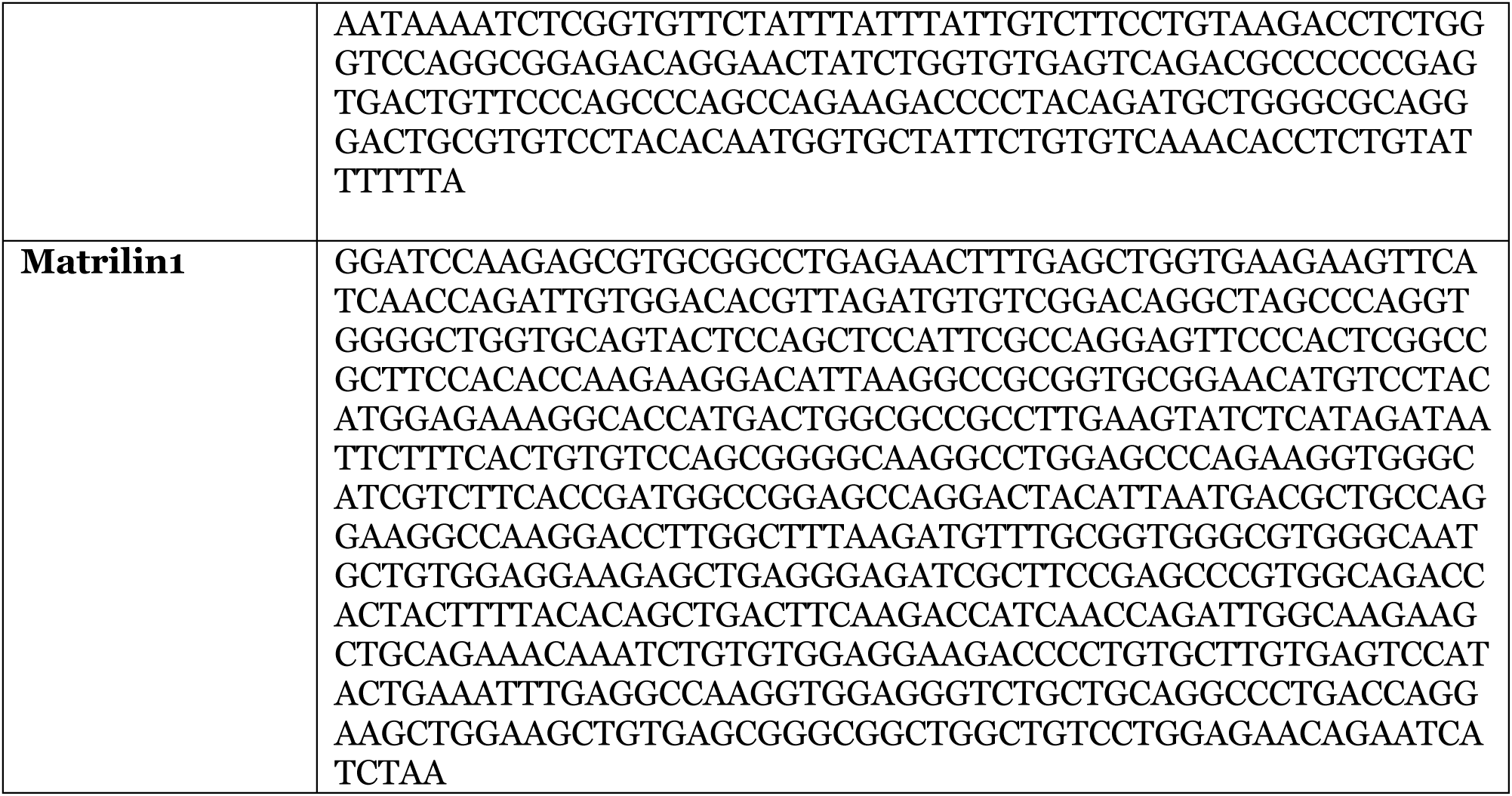
Target sequence (sense strand) of in situ hybridization probes.

**Supplementary Table 11:**
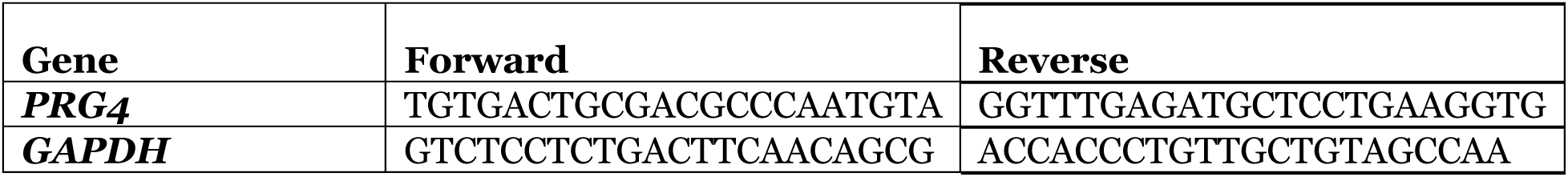
PCR primers for human *PRG4* and *GAPDH*.

